# Universal physical principles of protein structural genesis emerge in language-model representation space

**DOI:** 10.64898/2026.02.20.706798

**Authors:** Liu Chuanyang, Jiyuan Liu, Xinyuan Qiu, Xiaomin Wu, Wenying Li, Lu Min, Guohao Zhang, Shaowei Zhang, Lingyun Zhu

## Abstract

Protein structure is usually treated as the endpoint of sequence by most AI models, yet its true biological emergence is a process of ordered change, whose logic remains hidden in opaque black box. ***ProtGenesis*** creates a bidirectional mirror world: it maps amino acid assembly, elongation and mutation in biological space into quantitative **trajectories and ensembles** in protein language model representation space, then translates spatial geometry into testable biophysical hypotheses. Across peptides, reporter proteins and protein families, three universal general principles emerged: hierarchical directional assembly (**Principle I**), quantitative and reproducible structural-emergence trajectories (**Principle II**) and discrete topological transitions (**Principle III**) from short- to long-range order. Three novel metrics of spatial features [*D*, *ρ*, *δ*], complemented by squared Gaussian Wasserstein-2 measures, located structural anchors, sensitive regions and state transitions, enabling split-protein engineering and programmable protein design. ProtGenesis makes latent representations mechanistically interpretable, offering AI a route to discovering, rather than merely predicting, scientific principles.

**Graphical Abstract:** 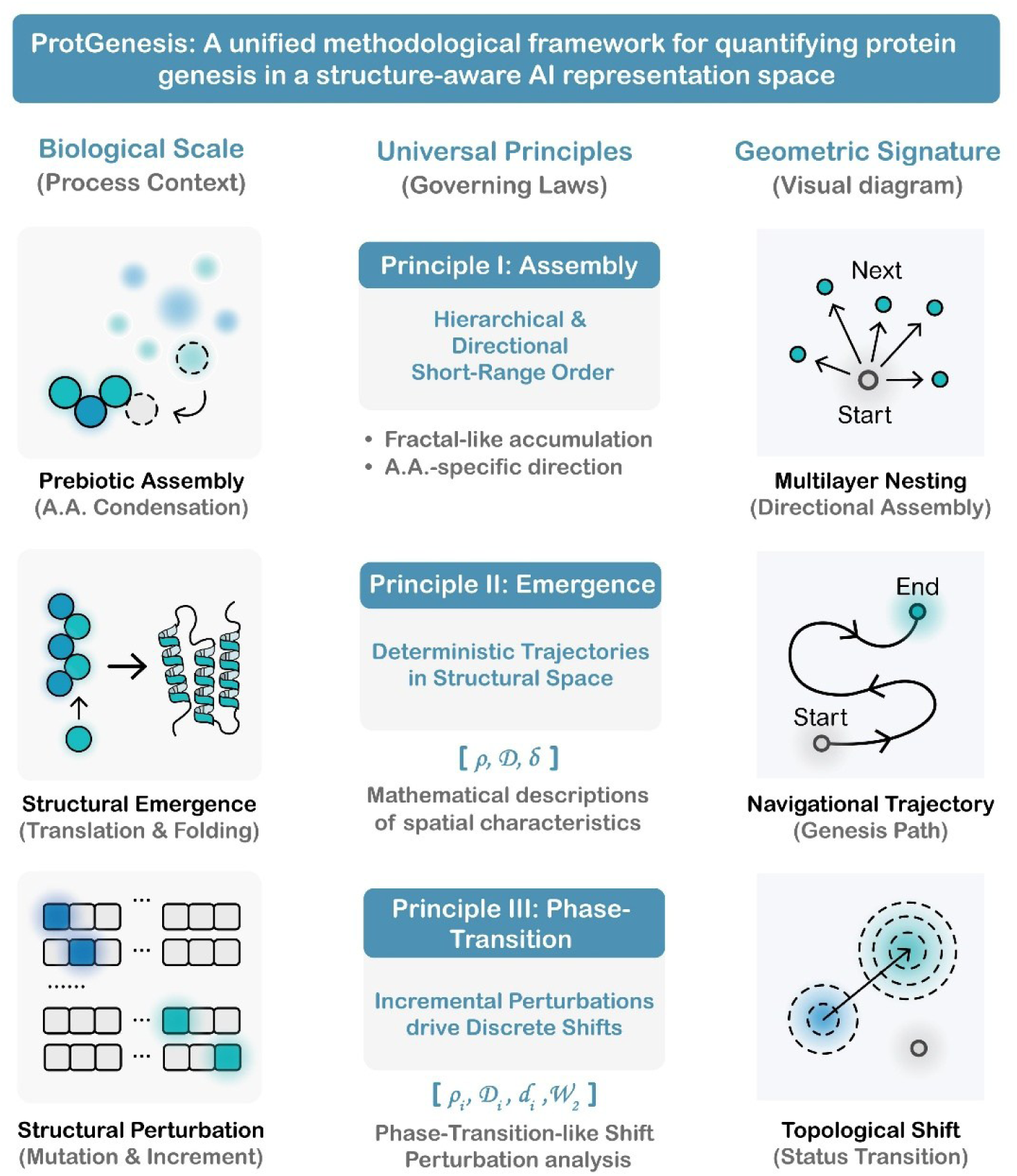

## Background

The origin of functional proteins stands as one of biology’s most profound enigmas, delineating the critical transition from prebiotic chemical chaos to the organized molecular architectures that sustain life^1–6^. While theoretical frameworks like the RNA world hypothesis^6,7^ and experimental reconstructions of prebiotic oligomerization by the Miller-Urey experiment^8^ have addressed the chemical origins of amino acids and other prebiotic building blocks, they primarily describe molecular composition rather than the determinants of structural causality. Anfinsen’s dogma posits that protein’s natural structure is only determined by its amino acid sequence, which provides a static, qualitative endpoint rather than a quantitative, dynamic process^9,10^. Thus, protein genesis remains conceptually fragmented across disparate scales, spanning prebiotic amino acid condensation, translation-coupled folding, and evolutionary diversification^11–13^. Crucially, the underlying physical laws governing the *process* itself, ***how discrete amino acid sequences deterministically navigate a vast structural space to achieve functional topology***, remain unresolved. Consequently, a unified quantitative framework capable of linking local molecular assembly rules to global structural emergence is missing, creating a fundamental disconnect between mechanistic biophysical understandings and biological reality.

Recent breakthroughs in artificial intelligence have further highlighted this gap. While deep learning models such as AlphaFold3^14^ and Boltz-2^15,16^ have enabled high-fidelity prediction of protein structures and biomolecular interactions and constructed comprehensive structural repositories (or protein universe) like the AlphaFold Protein Structure Database^17–25^, ***This prevailing single-structure-centric paradigm largely addresses “what structure” a sequence adopts, but not “how structure emerges”*** ^24,26–29^, leaves the intrinsic principles of genesis process of protein structure itself, and the mathematically organizing logic of the protein universe, uncharacterized^30^. Protein language models (PLMs, eg. ProstT5^31^, ProtT5^32^, VenusX^33^, and ESM3^34^) offer a transformative alternative by compressing proteins into high-dimensional numerical embeddings that implicitly encode fundamental biophysical constraints and evolutionary relationships^33,35–37^. Yet, these high-dimensional latent space are often treated as opaque computational **“black boxes”** rather than navigable physical spaces.

Here, we introduce ProtGenesis, a unified geometric framework that recasts protein genesis as a structured navigation of discrete sequence states through a continuous, high-dimensional representation space. We hypothesize that ***this space is not a stochastic continuum but a hierarchically organized physical system, quantitatively reflecting deterministic biological processes*** of protein genesis including amino acid assembly, translation and folding processes, and evolutionary adaptation. This perspective enables systematic exploration of protein genesis along biologically grounded trajectories, including stochastic prebiotic assembly of amino acids, stepwise residue addition mimicking translation, sequential substitutions reflecting evolutionary phylogenesis, and rational fixed-backbone protein design. Our work uncovers three organizing principles that collectively describe this process: **Hierarchical short-range assembly (*Principle I*)**, **Deterministic genesis paths (*Principle II*)**, and **Discrete topological phase-transition-like shifts (*Principle III*)**. Consequently, our work turns protein language model representation space into a measurable framework for analysing principle-governed structural organization, establishing an interpretable mathematical foundation that bridges prebiotic chemistry, evolutionary biology, and AI-driven *de novo* design

## Results

### 1. ProtGenesis: A unified methodological framework for quantifying protein genesis in a structure-aware representation space

To examine whether protein structural emergence can be analyzed as a measurable process while uncover the ***first principles*** of protein genesis process, we introduce ProtGenesis: a standardized and unified methodological process-aware framework that maps predefined biological sequence operations into a structure-aware protein language model representation space (***Fig. 1***). The framework converts amino acid assembly, sequence elongation, module integration and mutation into bottom-up, quantifiable trajectories, enabling systematic analysis from individual residues (zero) to fully folded protein architectures (one). **The core concept of ProtGenesis is a bidirectional mathematical mapping: *<u>biological sequence operations are encoded as calculable geometric objects in representation space, and the resulting geometry is used to formulate quantitative hypotheses about protein structural emergence</u>***.

**Fig. 1.**
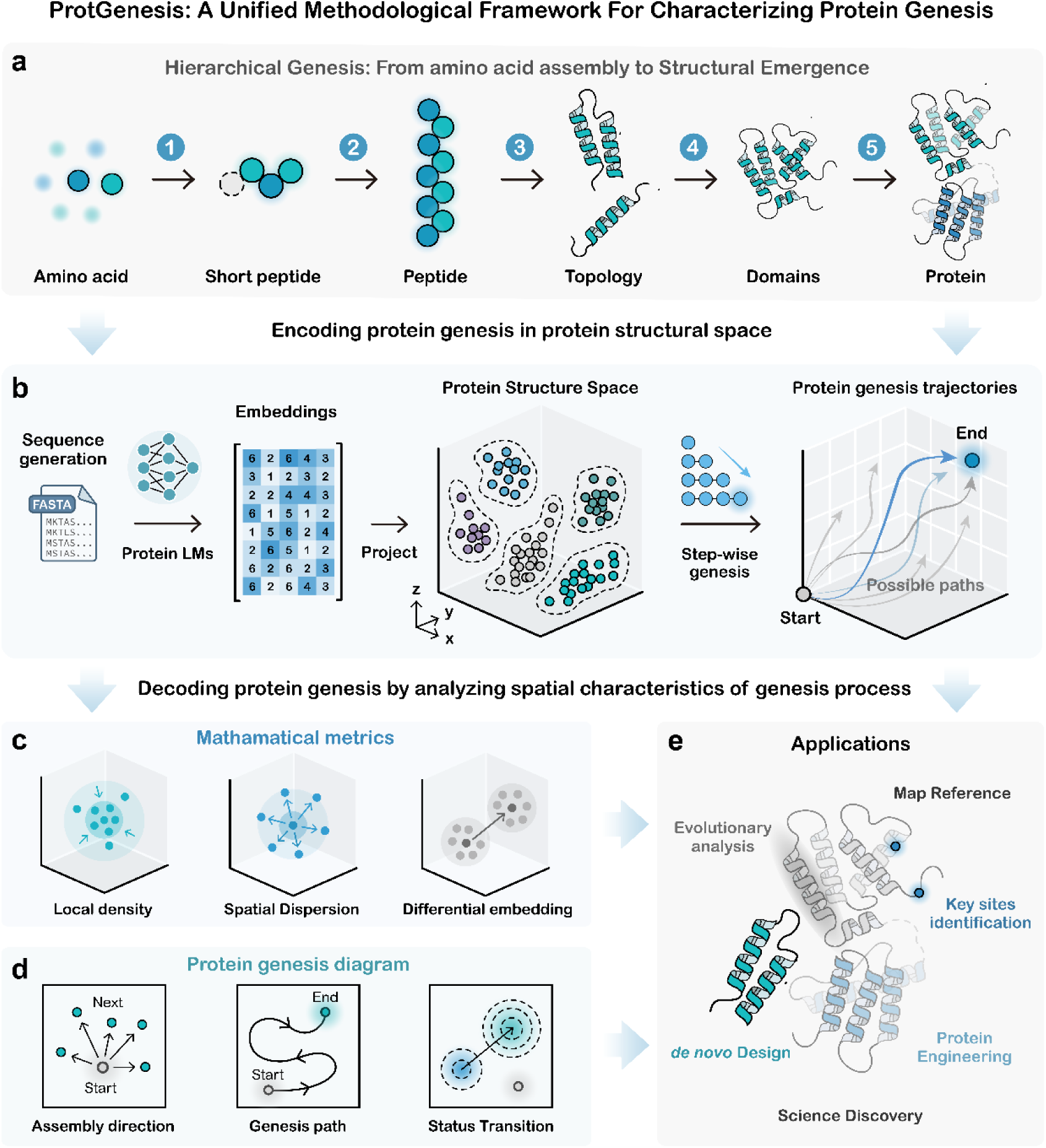
ProtGenesis: A unified methodological framework for quantitatively characterizing protein genesis in a structure-aware representation space. **a,** Hierarchical genesis of functional proteins is depicted as a structured, stepwise progression from amino acid assembly through peptides and topologies to domains and fully folded protein architectures. **b,** Biologically defined sequence series are generated, converted into high-dimensional embeddings and subjected to automated mathematical analysis using the ProtGenesis pipeline implemented in R and Python. **c,** Successive embedding states are treated as spatial trajectories and quantified by three metrics: Local Density (*ρ*), Spatial Dispersion (*D*), and Differential Embedding (*δ*). **d,** Three visualization modes, Assembly Direction Maps, Genesis Path Maps and Status Transition Maps, for representing directional organization, sequence-elongation trajectories and perturbation-associated state changes. **e,** ProtGenesis supports multiscale applications in protein-family analysis, structural landmark identification, split-protein engineering and protein design.

ProtGenesis integrates sequence construction, protein language model embedding, high-dimensional metric analysis and biological interpretation into a single workflow (***Fig. 1a–e***). By representing discrete sequence states within a continuous, structure-aware representation space and defining its trajectories mathematically, ProtGenesis provides a reproducible, data-driven foundation for uncovering the organizational principles of macromolecular assembly. Successive sequence states are encoded as high-dimensional representations and analyzed using Local Density (*ρ*), Spatial Dispersion (*D*) and Differential Embedding (*δ*), revealing spatial convergence, heterogeneity and abrupt transitions along protein genesis trajectories (***Fig. 1c***). The framework provides three complementary analytical views. Assembly Direction Maps describe residue-associated displacement directions, Genesis Path Maps trace the representation-space trajectories of sequence elongation, and Status Transition Maps describe distributional changes associated with module integration or mutation (***Fig. 1d***). Finally, the framework enables principle-driven applications across structural map referencing, evolutionary path analysis, critical-site identification, AI-guided protein design, engineering of split-protein systems and representation-space-based scientific discovery (***Fig. 1e***).

### 2. Principle I: Systematic peptide enumeration reveals context-sensitive hierarchical short-range order governed by directional assembly

To isolate the most primitive rules of protein genesis, we designed two complementary amino acid assembly regimes to probe short-range organization under distinct scenarios. First, to model minimal de novo amino acid condensation relevant to prebiotic peptide formation, we exhaustively enumerated all canonical peptide sequences of lengths 1-4, yielding 168,420 constructs (***Fig. 2a-c***; ***Supplementary Fig. S1***). Second, to model scaffold-constrained genesis through stepwise terminal extension, we enumerated all possible one- to four-residue N- and C-terminal extensions of a folded GFP scaffold, yielding 336,840 constructs (***Fig. 2d***; ***Supplementary Figs. S2*** and ***S3***). This paired design allowed us to determine how residue-associated assembly rules are retained and modulated across minimal and scaffold-constrained molecular contexts.

**Fig. 2.**
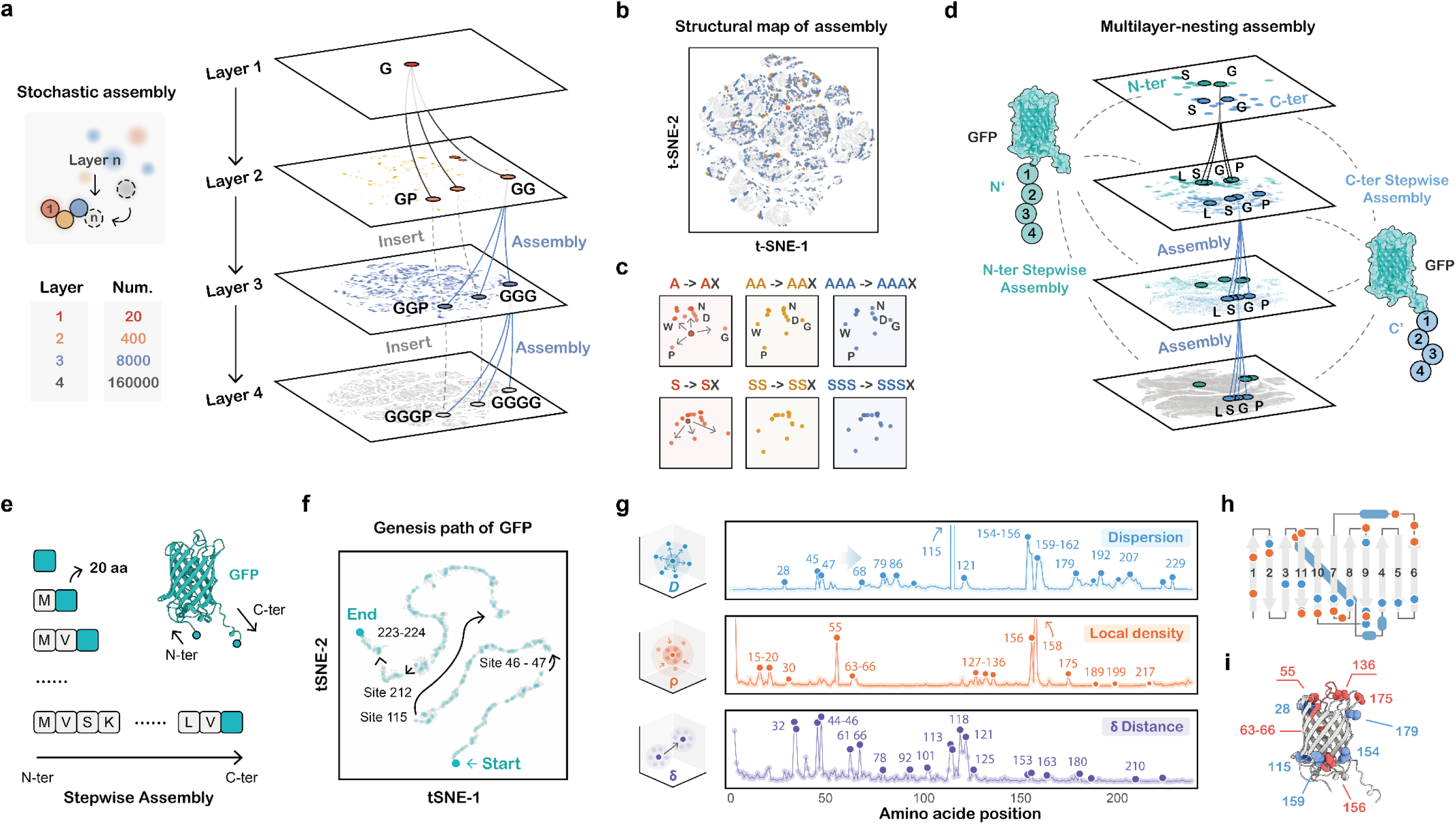
Universal principles of protein genesis: hierarchical directional assembly and reproducible emergence trajectories. **a-d**, The Assembly Principle (Principle I). **a,** Structure-aware representation-space maps derived from the systematic enumeration of 168,420 peptides of lengths 1-4, showing multilayer nesting. **b,** Hierarchical expansion across successive peptide layers. **c,** Assembly Direction Map revealing amino-acid-associated directional signatures. **d,** Organized representation-space patterns formed by N- and C-terminal GFP extensions. **e–i,** The Emergence Principle (Principle II). **e,** Schematic of stepwise GFP sequence elongation from successive N-terminal prefixes. **f,** Genesis Path Map showing the resulting representation-space trajectory. **g,** Quantitative decomposition of the trajectory using Local Density (*ρ*), Spatial Dispersion (*D*) and Differential Embedding Distance (*δ*). **h,i,** Mapping of metric-defined structural Fixed Points, Structural Pivots and structural Jumping Points onto GFP secondary and tertiary structures. Related assembly analyses are shown in ***Supplementary Figs. S1-S8***.

Mapping these ensembles into the structure-aware protein language model representation space via ProtGenesis revealed a multilayer nested architecture in which the *n* th layer was recursively organized around the (*n* − 1)th-layer foundation while expanding into previously unoccupied regions of representation space (***Fig. 2a***, ***b***; ***Supplementary Fig. S1a-c***). We quantified this organization using Differential Embedding (*δ*), which measures the representation-space displacement induced by residue addition. Across the short-peptide layers, context-extended transitions, such as AAA→AAAX, remained directionally aligned with their basal counterparts, such as A→AX, producing a residue-associated 20-direction assembly pattern visualized by the Assembly Direction Map and differential embedding heatmap (***Fig. 2c***; ***Supplementary Fig. S1c***). Although preceding sequence context progressively constrained the range of exploration, a shared residue-specific directional component remained prominent within the minimal peptide regime. These results indicate that short-range protein genesis follows an amino-acid-directed geometric logic.

Directional assembly signatures remained detectable on the full-length GFP scaffold, while their strength and orientation were quantitatively modulated by the surrounding sequence context (***Fig. 2d***; ***Supplementary Figs. S2***, ***S3*** and ***S7***). For a sequence context *c*, we defined the assembly vector associated with appending residue *X*as *v_c_*(*X*) = *E*(*cX*) − *E*(*c*), where *c* denotes the relevant precursor sequence. In the minimal de novo regime, *c*corresponds to one of the 20 single-residue contexts; in the GFP-scaffold regime, *c* denotes the corresponding scaffold precursor. In the minimal *de novo* regime, the 20 context-conditioned vectors showed high directional agreement, with a mean pairwise cosine similarity of 0.83 and a mean cosine similarity of 0.91 relative to their average vector. For GFP-scaffold extensions, the corresponding similarities were 0.43 and 0.67, respectively, and the mean vector magnitude was approximately 50-fold smaller, reflecting the dominant contribution of the full-length scaffold to the mean-pooled representation (***Supplementary Fig. S7a-c***). Despite this strong scaffold constraint, N- and C-terminal extensions formed organized, terminal-specific filling patterns within distinct representation-space regions (***Fig. 2d***; ***Supplementary Figs. S2*** and ***S3***). Across the protein language models examined, glycine consistently produced the largest mean *de novo* assembly-vector magnitude, further supporting a reproducible residue-associated component in the earliest assembly regime (***Supplementary Fig. S7d***).

This short-range organization extended into longer compositionally simple sequences, including (*G*)*_n_*, (*S*)*_n_* and other homopolymers, which followed structured, position-dependent trajectories as their chains elongated (***Supplementary Figs. S4*** and ***S5***). Along representative homopolymer trajectories, consecutive assembly-vector similarity changed sharply at the earliest elongation steps and subsequently stabilized at 0.94–0.97 for *n* ≥ 3, whereas vector magnitude progressively decayed toward a plateau (***Supplementary Figs. S5*** and ***S7***). This behavior indicates an early context-dependent reorientation followed by stable directional propagation under increasingly constrained sequence conditions. Flexible linkers, such as (*GGGGS*)*_n_*, which are widely used in protein engineering to preserve functional independence between fused domains, also exhibited ordered stepwise trajectories governed by the same interplay between residue-associated directionality and cumulative sequence constraint (***Supplementary Figs. S4*** and ***S5***). These observations place experimentally useful low-complexity motifs within a common quantitative assembly framework.

To determine whether these directional assembly patterns extend to amino acid compositions associated with the earliest stages of protein evolution, we examined homopolymers composed of prebiotically relevant amino acids, including poly-G, poly-A, poly-D, poly-V and poly-E^7,8,11^. Projecting these sequences into the ProtGenesis space revealed distinct, amino-acid-associated propagation trajectories from the earliest stages of chain extension (***Supplementary Fig. S6a-d***). At longer sequence scales, however, the accessible ensemble progressively converged: Spatial Dispersion *D* decreased strongly with genesis step in both combinatorial early-amino-acid peptides (*r_s_* = −0.986, *P* = 2 × 10^−23^) and homopolymers (*r_s_* = −0.919, *P* = 2 × 10^−49^; ***Supplementary Fig. S8a-c***). Together with the accompanying density and differential-distance profiles, these results reveal a scale-dependent transition from pronounced residue-directed exploration at short lengths to context-guided spatial convergence during extended genesis (***Supplementary Fig. S8***).

Collectively, these results establish the **<u>Assembly Principle</u>** (**<u>Principle I</u>**): *Protein genesis is governed by hierarchical short-range order arising from directional amino acid assembly*. Amino acid identity defines the initial direction of representation-space propagation, whereas increasing sequence length and scaffold context progressively channel, attenuate and stabilize that direction. The Assembly Principle therefore unifies intrinsic residue-associated directionality with context-dependent convergence, providing a multiscale assembly logic that connects minimal peptide formation, low-complexity sequence extension and scaffold-constrained protein growth (*Fig. 2a-d*; *Supplementary Figs. S1-S8*).

### 3. Principle II: Growing proteins trace reproducible representation-space trajectories associated with higher-order organization

To quantify how long-range order emerges from short-range rules, we systematically simulated the residue-by-residue genesis of GFP by extending the sequence from the N- to the C- terminus (***Fig. 2e***, ***f***). This computational sequence-elongation model provides a representation-space analogue of co-translational protein emergence. GFP genesis traced a directed, reproducible trajectory through the structure-aware protein language model representation space, indicating that higher-order organization emerges through an oriented spatial program (***Fig. 2f***). To decode this trajectory, we extended *δ* into a tripartite spatial metric system comprising Local Density (*ρ*), Spatial Dispersion (*D*) and Differential Embedding (*δ*), which resolves the sequence-elongation path into discrete, biologically meaningful transition events (***Fig. 2g***). Differential Embedding quantified sequential displacement, cosine distance captured directional reorientation where indicated, and Spatial Dispersion and Local Density characterized the spread and compactness of local sequence ensembles (***Fig. 2g***; ***Supplementary Fig. S10***). Mapping these metric-defined events onto the AlphaFold3-predicted GFP structure identified three classes of critical residues with distinct structural roles (***Fig. 2h***, ***i***).

First, peaks in *ρ* identify structural “***Fixed Points***” (for example, positions 55, 136 and 175), which function as structural anchors. These points consistently localize to loop regions or topological initiation and termination sites (***Fig. 2h***, middle panel). Analytically, residue *j^ρ^* marks the establishment of a dense structural core, whereas *j^ρ^* + 1 serves as a quantitative marker for the initiation of a new stable subdomain or secondary-structure element. Second, surges in *D* reveal “***Structural Pivots***” (for example, positions 28, 115 and 154). These high-dispersion nodes coincide with intramolecular interaction interfaces and cooperativity centers critical for structural integrity (***Fig. 2i***). They represent susceptible sites at which conformation is highly sensitive to sequence perturbation. Consequently, mutational analysis should consider *j^D^* together with its immediate neighbors, *j^D^* − 1 and *j^D^* + 1, because substitutions within this window are most likely to trigger substantial structural reorganization. Third, spikes in *δ* mark structural “***Jumping Points***” (for example, positions 66, 101 and 118). These discontinuities signify discrete topological transitions associated with the nucleation of secondary structures or domain closure, indicating that genesis proceeds through punctuated consolidation (***Fig. 2g-i***). For engineering purposes, residues at *j^δ^* and *j^δ^* + 1 define the critical step at which the system transitions between distinct structural states.

Collectively, the concordance between these spatial metrics and structural landmarks establishes the **<u>Emergence Principle</u>** (**<u>Principle II</u>**): *Functional protein genesis is not a continuous drift but follows a directed, reproducible trajectory* punctuated by structural Fixed Points, Structural Pivots and structural Jumping Points. The tripartite metric system thereby transforms the abstract concept of structural emergence into a measurable representation-space process defined by identifiable spatial coordinates.

### 4. Principle III: Incremental sequence alterations drive discrete topological phase transition-like shift from local to global

**Principles I** and **II** establish that protein genesis proceeds through hierarchical assembly along directed trajectories, functional folding further requires a transition from local short-range order to global topological coherence. We therefore posited that ***continuous sequence change, such as stepwise genesis and cumulative mutation, can trigger discontinuous structural reorganization*** and tested this hypothesis at two coupled scales: <u>a macroscopic regime</u> in which residue accrual is coarse-grained into module integration, and <u>a microscopic regime</u> in which single-residue substitutions generate mutational ensembles reporting state stability and switching.

<u>At the macroscopic scale</u>, we examined whether the continuous GFP genesis path can be resolved into discrete representation-space states. Using AlphaFold3 topology, we decomposed the 239-residue GFP into 11 structurally coherent modules (***Fig. 3a-c*** and ***Supplementary Fig. S9a***) and reconstructed genesis using four distinct assembly strategies. Across all strategies, representation-space trajectories exhibited extended periods of gradual integration punctuated by abrupt, reproducible displacements that recurred across pathways at the module scale (***Fig. 3b***, ***c***). In Pathway 1, incorporating fragments spanning modules 3–5, which complete major elements of the β-barrel, induced a pronounced reorganization toward global coherence. The N-to-C trajectory showed cosine-based *δ* peaks at F2 and F4, whereas fragment dispersion *D* reached its maximum at the F4 accumulated fragment (*D* = 2.93), which contains the chromophore-containing module; local density *ρ* decreased monotonically across the assembly sequence. Together, these metrics localized the dominant transition to chromophore-pocket formation (***Supplementary Fig. S10a-c, Supplementary Table 1***). In Pathway 2, a discontinuity at the 7→8→5 module-assembly step marked secondary-structure closure. AlphaFold3 models placed these representation-space transitions in the context of domain nucleation and packing convergence (***Fig. 3c*** and ***Supplementary Fig. S9b***), consistent with the emergence of long-range organization through discrete fold-associated transitions.

**Fig. 3.**
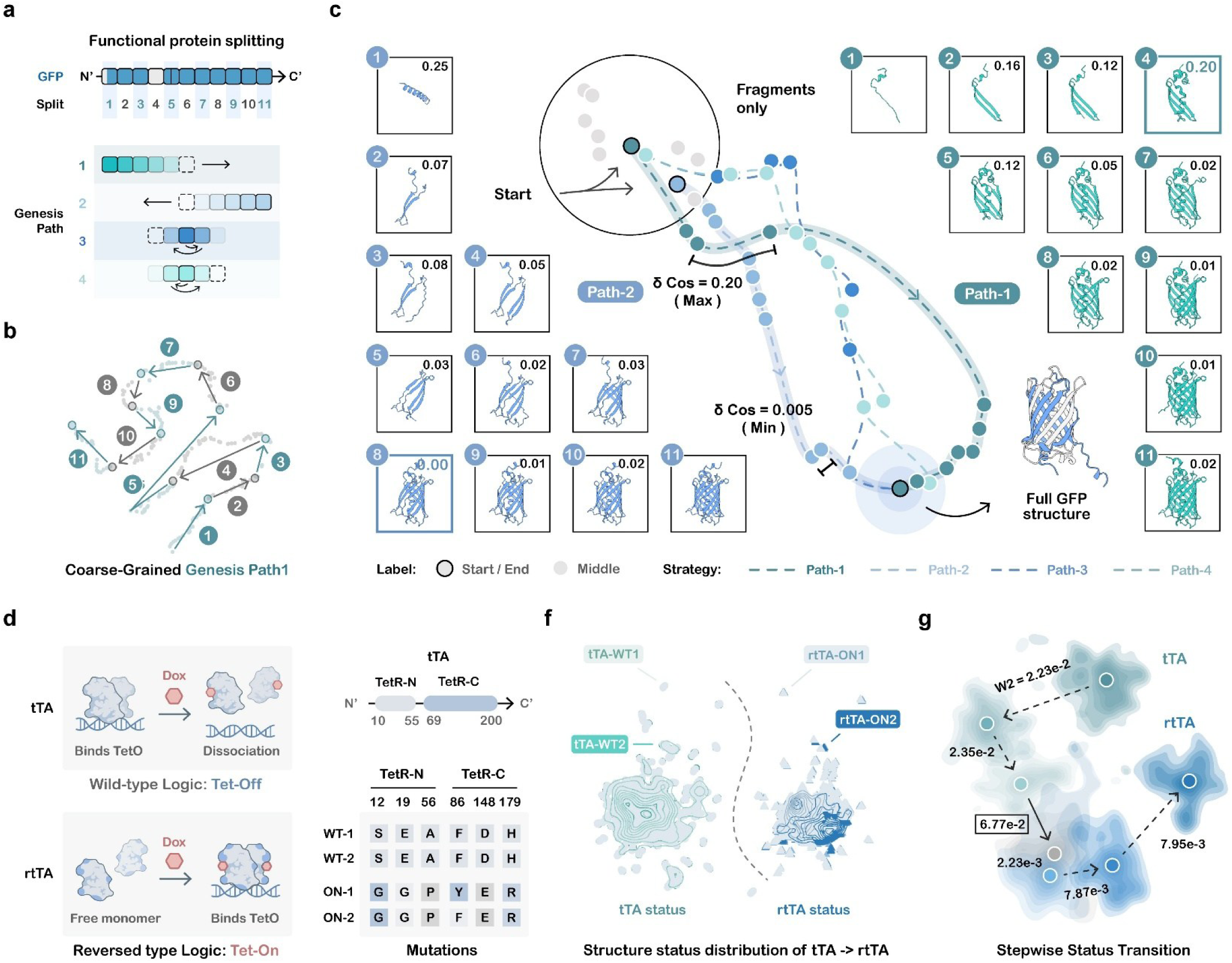
Principle III: Incremental sequence alterations drive discrete topological state transitions at macroscopic and microscopic scales. **a-c,** Macroscopic coarse-graining of GFP genesis. **a,** Decomposition of GFP into 11 structurally coherent topological modules based on the AlphaFold3-predicted structure. **b,** Four module-assembly strategies, N-to-C, C-to-N, midpoint-outward and midpoint-asymmetric, used to reconstruct global topology from local modules. **c,** Representation-space trajectories and metric profiles revealing quantized topological shifts, with the dominant N-to-C transition localized to the chromophore-containing F4 module. **d–g,** Microscopic state transitions associated with functional evolution. d, Stepwise substitution path converting Tet-Off tTA into Tet-On rtTA. **e,f,** Representation-space distributions of tTA- and rtTA-associated sequence states. **g,** Status Transition Map identifying A56P as the largest stepwise distributional change. Related analyses are shown in *Supplementary Figs. S9-S17*.

<u>At the microscopic scale</u>, we asked whether single-residue perturbations reveal transition boundaries analogous to those observed during genesis. Comprehensive single-site mutational enumeration of GFP produced bounded, position-specific distributions with interwoven geometries, in marked contrast to the quasi-linear sequence-elongation trajectory (***Supplementary Fig. S11a-d***). Each residue therefore defines a distinct, locally constrained representation-space neighborhood. This organization delineates the micro-topology of the mutational landscape, including accessible mutational paths and potential state-transition boundaries.

To quantify this micro-perturbative regime, we adapted our metrics into a mutation-indexed framework: Local Density (*ρ_i_*) and Spatial Dispersion (*D_i_*) characterize the ensemble geometry at each site, whereas Differential Embedding Distance (*δ_i_*) measures the displacement from the wild-type state. These metrics revealed that *ρ_i_* and *D_i_* are sharply modulated by fold topology, identifying structural anchors and susceptibility nodes (***Supplementary Fig. S11e***).

Together, they define the local roughness of the mutational landscape. Cross-model analysis further identified residue 83, located in a turn, in four of five models and residue 221, located in β11, in three of five models (***Supplementary Figs. S12*** and ***S13***). Both residues were fully conserved among 13 GFP homologs and ranked at the 100th percentile of the position-wise conservation distribution. The N-to-C genesis-associated Jumping Points at residues 66, 101 and 118 showed lower conservation, with residue 66 ranking at the 4.2nd percentile, indicating that mutational susceptibility and elongation-associated transitions capture distinct dimensions of structural organization (***Supplementary Fig. S13***). A length-matched random-sequence control yielded uniformly low-confidence structural predictions, with a mean normalized pLDDT of 0.413 ± 0.009 and all values below 0.7. Its representation-space dispersion and density (*D* = 0.89, *ρ* = 0.58) contrasted sharply with those of the GFP ensemble (*D* = 0.16, *ρ* = 0.93), supporting the fold-associated specificity of the GFP mutational geometry (***Supplementary Fig. S14***).

Crucially, to directly link micro-geometry to functional switching, we analyzed the Tet-ON/OFF system, in which stepwise mutations convert the DNA-binding phenotype of tTA into the reverse rtTA phenotype (***Fig. 3d***). The Status Transition Map showed that tTA- and rtTA-associated sequence states occupy distinct, non-overlapping representation-space basins (***Fig. 3e***, ***f***). The trajectory remained within the tTA-like basin until the A56P substitution, at which point it underwent an abrupt transition into the rtTA-like state (***Fig. 3g***).

Because a phase-transition-like event concerns a change in the distribution of sequence states, we quantified each adjacent state pair using the squared Gaussian Wasserstein-2 distance (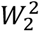). For each protein language model, the raw embedding ensembles were independently projected into a model-specific 32-dimensional random Gaussian projection space generated with seed 42, and 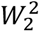 was calculated from the mean and regularized covariance of each projected state ensemble. Manhattan, Euclidean and cosine measures provided complementary descriptions of single-step displacement and directional reorientation.

Four protein language models, ProstT5, ProtT5, ESMC-6B and METL-G^38^, consistently identified the MutAdd_2_→MutAdd_3_ step corresponding to A56P as the largest distributional shift (***Supplementary Fig. S15*** and ***Supplementary Table 2***). Because the raw values are model-scale-specific, cross-model comparison was based on within-model standardized scores.

In ProstT5, A56P was also maximal under four complementary measures: squared Gaussian Wasserstein-2 distance (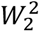), mean *L*_2_ displacement, mean *L*_1_ displacement and cosine displacement (***Supplementary Fig. S15c-f***). Single-mutation ensembles comprising 3,915 sequences per state sharply resolved consecutive states and localized the dominant transition to A56P (***Supplementary Fig. S16***). A complementary BioEmu^39^-based conformation ensemble simulation analysis of 500 conformations for each endpoint produced overlapping conformational ensembles, revealing distinct resolutions of sequence-representation and conformational landscapes (***Supplementary Fig. S17***). A56P is the established rtTA2s-M2 mutation, independently recovering a known functional-switching substitution as the dominant representation-space transition. The recurrence of the A56P-associated transition across four protein language models, together with the cross-model agreement of GFP mutation-associated profiles, separates model-shared geometric signals from model-specific representation features.

Together, these analyses establish the **<u>Phase-Transition Principle</u>** (**<u>Principle III</u>**): *Incremental sequence alterations, arising through ordered module accrual or single-residue mutation, generate gradual progression punctuated by discrete topological state transitions*. These transitions partition representation space into quasi-stable regimes separated by critical boundaries, providing a unified quantitative description of structural assembly, folding and functional diversification (***Fig. 3a-g*; *Supplementary Figs. S9-S17***).

### 5. Emergence-associated representation features recur in TRIM-family proteins

To test whether these emergence rules generalize beyond model systems, we analyzed the stepwise genesis of two human TRIM-family E3 ligases, TRIM21 and TRIM5^40–42^. Despite diverging to support distinct innate immunity functions, both proteins followed highly structured genesis trajectories and shared representation-space signatures (***Supplementary Fig. 18***). Recurring density-defined anchors and dispersion-defined susceptibility nodes coincided with modular boundaries in AlphaFold3-predicted structures (***Supplementary Fig. 18a∼d***). Notably, local density surges identified structural ***Fixed Points*** within the early RING-domain region (residues ∼16–56), corresponding to the emergence of the catalytic E3 ligase core as a discrete folding module (***Supplementary Fig. 18a***, ***9b***). Furthermore, metric-defined transition sites aligned with known structure-function determinants: In TRIM21, signatures at position 383 corresponded to the B30.2/PRY domain’s hydrophobic pocket (involving residues such as W383 and F450), while dispersion-enriched sites around position 180 aligned with coiled-coil dimerization motifs critical for higher-order assembly (***Supplementary Fig. 18a, b***). Analogous density and dispersion patterns recurred in TRIM5 at corresponding architectural stages (***Supplementary Fig. 18c, d***), indicating that ***some representation-space features may recur across related proteins with modular fold organization despite substantial sequence divergence*.**

Together, these observations provide an initial cross-protein consistency test of the Emergence Principle (Principle II) and support the use of density- and dispersion-associated representation-space features as candidate modular and functional landmarks across related TRIM proteins.

### 6. Representation-space fixed points nominate candidate split-protein sites

To examine the engineering utility of representation-space features, we hypothesized that the signatures identified by our metrics, specifically ***Structural Fixed Points*** defined as peaks in *ρ*, could provide structure-guided coordinates for prioritizing candidate split-protein cleavage sites. A central challenge in split-protein design is identifying permissive cleavage sites that yield folding-competent fragments. We reasoned that Structural Fixed Points could mark structurally permissive modular boundaries for protein splitting.

To validate this, we systematically analyzed the local density profiles of four diverse proteins: GFP, the tetracycline repressor (TetR), Cre recombinase and HaloTag enzyme^43^. In GFP, established split sites (residues 157 and 214) aligned precisely with prominent density peaks (***Supplementary Fig. 19a***, ***19b***). These topological anchors also matched sites identified by ***ProDomino***^44^, a state-of-the-art machine learning pipeline for predicting protein domain-insertion sites, suggesting that representation-space compactness may capture aspects of modular organization recognized by other computational approaches.

We then expanded the analysis to functional effectors. In TetR, the validated split site at residue 99 corresponded to a major density maximum at the junction between the DNA-binding and dimerization domains (***Fig. 4a, b***). Similarly, in Cre recombinase and HaloTag, literature-reported split sites aligned with Fixed Points identified by our framework (***Fig. 4c, d***; ***Supplementary Fig. S19c, d***). This correspondence supports the use of representation-space features for candidate-site prioritization.

**Fig. 4.**
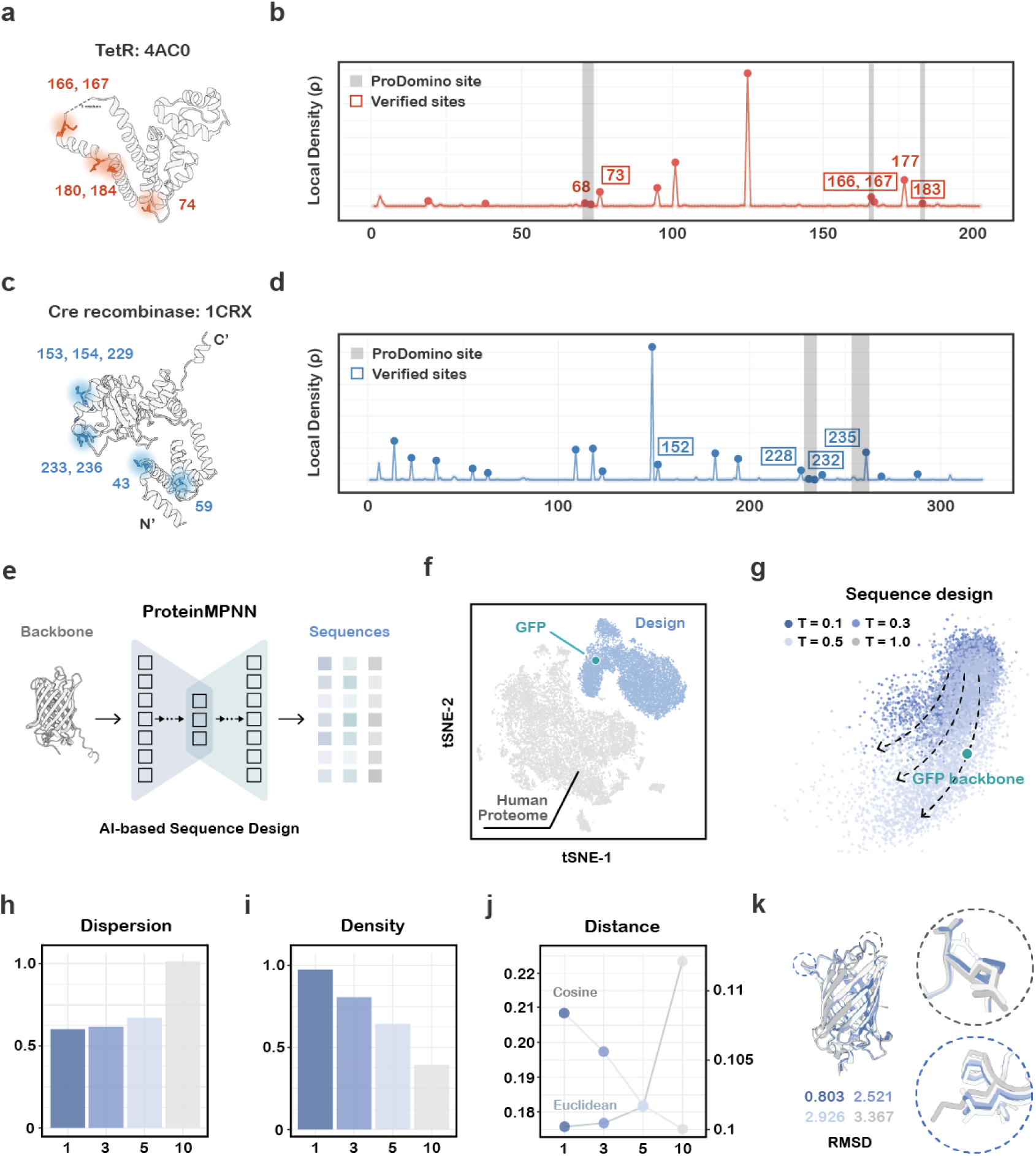
Representation-space coordinates for rational protein engineering and fixed-backbone design. **a,b,** AlphaFold3-predicted structure of TetR and its local-density profile, showing the reported split-protein site and candidate density-associated Fixed Points. **c,d,** Corresponding structural mapping and local-density profile for Cre recombinase; shaded regions indicate sites identified by ProDomino where applicable. **e-k,** ProteinMPNN-based exploration of fixed-backbone GFP design space. **e,** Generation of fixed-backbone sequences at controlled sampling temperatures. **f,** Distribution of 8,000 designed variants relative to the human-proteome background. **g,** Principal component analysis visualization of the temperature-associated design trajectory. **h-j,** High-dimensional characterization using Spatial Dispersion (*D*), Local Density (*ρ*)and Differential Embedding Distance (*δ*). **k,** AlphaFold3-predicted structures of representative designed variants. Related analyses are shown in ***Supplementary Figs. S19*** and ***S20***.

By quantifying these intrinsic boundaries, ProtGenesis provides a quantitative, principle-guided basis for prioritizing candidate cleavage sites and modular boundaries before experimental screening.

### 7. Sampling temperature enables programmable exploration of fixed-backbone protein design space

Deep learning models such as ProteinMPNN^45^, RFdiffusion^46^ have revolutionized protein engineering through *de novo* protein design. We next asked whether ProtGenesis could quantify how a generative protein-design model explores sequence space. Using ProteinMPNN with GFP as a fixed backbone, we generated 2,000 designed sequences at each of four sampling temperatures, yielding 8,000 fixed-backbone variants in total (***Fig. 4e***; ***Supplementary Fig. S20a***). With the GFP backbone fixed throughout, we refer to these sequences as fixed-backbone designed variants.

Mapping these designs revealed that they occupied a representation-space region distinct from the human-proteome background (***Fig. 4f***). Increasing the sampling temperature *T* drove the design ensemble along a directional trajectory away from the native GFP state, revealing an organized structural flow through representation space (***Fig. 4g***; ***Supplementary Fig. S20a***). This “Structural flow” indicates that *T* acts as a tunable control parameter for steering exploration toward new representation-space regions. Our quantitative framework captured this programmable exploration. As *T* increased, *δ* and *D* scaled monotonically, reflecting a controlled expansion into new structural territories (***Fig. 4h–j***; ***Supplementary Fig. S20b, c***). AlphaFold3 predictions supported this trend, showing that higher-temperature variants progressively deviated from the GFP fold while retaining core topology (***Fig. 4k***).

This analysis also highlighted the value of high-dimensional representation-space analysis. Low-dimensional projections compressed the design ensembles and obscured underlying trends, whereas the intrinsic high-dimensional metrics (*ρ*, *D*, *δ*) resolved the relationship between sampling temperature and structural displacement (***Fig. 4g-j***; ***Supplementary Fig. S20a-c***), underscoring the importance of high-dimensional analysis for protein design.

Collectively, these results show that ***fixed-backbone protein design follows a navigable trajectory through representation space***. ProtGenesis provides a quantitative coordinate system for monitoring and steering this exploration, turning generative design into a measurable, principle-guided process.

***ProtGenesis*** maps protein assembly and perturbation into quantitative trajectories in protein language model representation space. Across minimal peptides, growing proteins, mutational landscapes, natural protein families and fixed-backbone design ensembles, it reveals three recurring organizational features: context-sensitive hierarchical assembly, structured trajectories of structural emergence and discrete representation-space state transitions (***Figs. 2-4***; ***Supplementary Figs. S1-S20***). By linking biological operations to latent-space geometry and converting geometric signatures into testable structural hypotheses, ***ProtGenesis provides a new route to protein language model interpretability and a quantitative basis for candidate protein engineering strategies*.**

## Discussion

Here, we present ProtGenesis as a process-aware framework for interrogating protein language model representations. ProtGenesis represents each protein together with the sequence operations that generate and perturb it, including amino acid addition, sequence elongation, module integration and mutation. These operations are encoded as trajectories or local ensembles in a structure-aware representation space, and their geometry is translated into testable hypotheses about protein structural emergence. The central conceptual contribution is **bidirectional**: ***biological operations probe the model, while model geometry guides biological interpretation and experimental hypothesis generation***.

Across minimal peptides, growing proteins, mutational landscapes and natural protein families, three recurring organizing principles emerged: ***<u>hierarchical directional assembly, reproducible</u> <u>structural emergence and phase-transition-like state shifts</u>***. These principles explain the physical basis of **“more is different”** ---how functional order emerges from local rules through constrained transition paths, and indicate that ***functional structure is defined not only by its endpoint, but also by the ordered and discrete sequence of transitions through which that endpoint becomes reachable*.**

Notably, this underlying logic parallels the algorithmic determinism of ***cellular automata***^47^, in which macroscopic complexity emerges spontaneously through the iterative application of simple, local interaction rules among elementary units. Similarly, protein genesis can be viewed as a form of ***biological computation***^48^, in which discrete amino acid inputs drive the system through an intrinsically constrained structural program that channels molecular exploration toward higher-order organization. ProtGenesis makes this process mathematically measurable through a process-matched metric system. <u>Differential Embedding</u> (***δ***) canonically evaluated using the Manhattan norm, quantifies displacement between sequential states, <u>cosine distance</u> captures directional reorientation. <u>Spatial Dispersion</u> (***D***) and Local Density (***ρ***) characterize the spread and compactness of local ensembles; and the squared Gaussian Wasserstein-2 distance (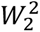) distinguishes distributional state shifts by integrating changes in ensemble mean and covariance. The convergence of complementary metrics, together with cross-model validation, ***identifies geometric events shared across protein language models while distinguishing model-specific representation features***. This framework enables applications ranging from rational domain definition and split-protein engineering to evolutionary analysis and the programmable steering of protein-design pipelines.

Beyond biological engineering, the framework offers a process-grounded route to AI interpretability. Through this **“Mirror-world” methodology**, biological processes are mirrored into latent representation space, and the mathematical organization of that space is translated back into biologically meaningful and experimentally testable hypotheses. We show that protein language models implicitly encode organizing principles of structural genesis, bridging data-driven deep learning with mechanistic scientific understanding. Consequently, ProtGenesis suggests a transferable paradigm for **AI for Science**: the same bottom-up assembly– perturbation logic may be used to interrogate the latent representations of other complex systems and extract candidate scientific principles from their geometry. This establishes a bidirectional paradigm in which ***AI serves as a computational laboratory for scientific discovery, while physical and biological principles reciprocally guide the design, interpretation and validation of AI models***.

In summary, we provide a unifying mathematical framework for protein structural genesis, spanning minimal peptide assembly, functional protein emergence, evolutionary perturbation and AI-guided protein design. ProtGenesis establishes a geometry-based and biologically grounded framework for biological engineering and offers a blueprint for interpretable, science-driven artificial intelligence across disciplines. More broadly, it advances a view of artificial intelligence beyond endpoint prediction: scientifically defined processes can be projected into latent spaces, mathematically interrogated and reflected back into the physical world as testable principles of biological organization.

## Methods and Material

### Combinatorial Peptide Generation and Dataset Construction

To reconstruct the fundamental organization of protein sequence-associated representation space from first principles, we performed a comprehensive combinatorial enumeration of peptide sequences. Using a custom R function, generate_stepwise_for_protein(), we hierarchically assembled all possible canonical amino acid sequences of lengths 1–4, yielding 20^1^ + 20^2^ + 20^3^ + 20^4^ = 168,420 unique peptides. To examine structural emergence in the context of a folded scaffold, we generated a comprehensive library of GFP fusions containing all possible one-to four-residue N-terminal and C-terminal extensions, yielding 336,840 unique constructs. Furthermore, to investigate assembly trajectories beyond the short-peptide regime, we constructed homopolymeric sequences, including poly-glycine, poly-serine and polyalanine, together with flexible linkers such as (GGGGS)*_n_*, with lengths extending to 120 residues. These sequence collections enabled direct comparison of minimal residue-associated organization, scaffold-dependent constraints and length-dependent changes in representation-space geometry. All sequences were stored in standardized FASTA format with unique identifiers for high-throughput computational processing.

For the early-Earth-motivated sequence analysis, we assembled three peptide cohorts. The EAC cohort comprised 100 combinatorial sequences generated from an early-amino-acid set and tracked over 30 genesis steps. The MOTIF cohort comprised three short sequence motifs— ferredoxin, P-loop and poly-glycine—tracked over seven steps. The POLY cohort comprised five homopolymers, poly-G, poly-A, poly-D, poly-V and poly-E, tracked over 120 steps. For each cohort, we calculated Spatial Dispersion *D* and Local Density *ρ* at every genesis step and evaluated their monotonic relationships with genesis step using Spearman correlation (***Supplementary Figs. S6*** and ***S8***).

### Protein Language Models and Embedding Generation

Quantitative structural descriptions were generated using ***ProstT5***^35^, a structure-aware protein language model. For each sequence, we extracted the 1,024-dimensional residue-level encoder representation and applied mean pooling across residues to obtain one fixed-dimensional vector per sequence. All primary distances and geometric metrics were calculated in the original high-dimensional representation space. Principal component analysis, t-distributed stochastic neighbor embedding and uniform manifold approximation and projection were used for visualization only.

For longer sequences, including GFP, TRIM21 and TRIM5, we applied the same embedding-extraction and mean-pooling procedure to every intermediate sequence state, thereby maintaining methodological consistency across sequence lengths. For the standard *D*, *ρ* and *δ* analyses, embedding matrices were mean-centred feature-wise and were not feature-scaled. The *W*_2_ analysis followed a separate model-specific branch in which raw mean-pooled embeddings were projected without centering or scaling, as described in the Distributional Transition Analysis section.

For cross-model validation, we additionally generated mean-pooled representations using ProtT5, ESMC-6B^49^, ESM2-650M and METL-G^38^ with the same residue-level pooling strategy. The complete GFP mutation analysis used five models, ProstT5, ProtT5, ESMC-6B, ESM2-650M and METL-G, whereas the tTA-to-rtTA transition analysis used ProstT5, ProtT5, ESMC-6B and METL-G. Because the models differ in embedding dimensionality and scale, model-specific metric profiles were compared using position-wise Spearman correlation, and distributional transition scores were standardized within model before cross-model comparison.

Mean pooling generated exact duplicate vectors for a subset of distinct GFP single-site mutants. These duplicate rows were quantified and reported together with the corresponding density analyses (***Supplementary Fig. S13***).

### Integrated Chomputational Framework and Visualization

***ProtGenesis*** was developed as an integrated R/Python computational framework combining sequence construction, embedding generation, high-dimensional matrix analysis, metric calculation and visualization. To visualize the organization of the high-dimensional representation space, PCA was used to display global variance and trajectory directionality, t-SNE was used to examine local neighborhood organization, and UMAP was used to visualize large sequence collections. Except for the Gaussian Wasserstein-2 analysis, all primary distances and geometric metrics (*D*, *ρ* and *δ*) were calculated in the original high-dimensional embedding space. Principal component analysis, t-distributed stochastic neighbour embedding and uniform manifold approximation and projection were used for visualization only.

The framework includes three complementary spatial metrics: Spatial Dispersion ***D***, Local Density ***ρ*** and Differential Embedding ***δ***. These metrics quantify the spread of a local sequence ensemble, the compactness of its local neighbourhood and the displacement between sequential sequence states, respectively. Assembly Direction Maps display residue-associated displacement vectors, Genesis Path Maps display sequence-elongation trajectories, and Status Transition Maps display distributional changes associated with module integration or mutation (***Fig. 1***, ***Figs. 2*** and ***3***).

The metric assigned to each analysis was matched to the biological object under investigation. *L*_1_ -based Differential Embedding Distance quantified sequential state displacement; cosine similarity and cosine distance quantified directional consistency and reorientation; Spatial Dispersion and Local Density described local ensemble variability and compactness; and Gaussian Wasserstein-2 distance quantified distributional shifts between ensemble states. This combination of metrics enabled the separation of directional movement, local convergence, ensemble variability and state transition within a unified analytical framework.

### Differential Embedding Analysis

To quantify the displacement associated with sequential sequence changes, we defined the Differential Embedding between a sequence state at hierarchical level *n* and its ancestral state at level *n* − *i* as

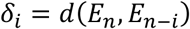

where *E_n_* and *E_n_*_−*i*_ denote the corresponding mean-pooled embeddings. The canonical Differential Embedding Distance used the Manhattan distance:

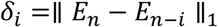

Here, *i* = 1 measures the immediate displacement caused by one sequence operation, whereas *i* > 1 captures longer-range coupling between sequence states. Euclidean and cosine distances were used as complementary sensitivity measures where indicated.

For a sequence context *c*, we defined the assembly vector associated with appending residue *X*as *v_c_*(*X*) = *E*(*cX*) − *E*(*c*), where *c* denotes the relevant precursor sequence. In the minimal de novo regime, *c*corresponds to one of the 20 single-residue contexts; in the GFP-scaffold regime, *c*denotes the corresponding scaffold precursor.

For a homopolymer precursor *H_n_*, the assembly vector was defined as

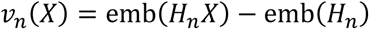

with *H_n_* denoting the corresponding *n*-residue precursor. Consecutive direction similarity was calculated as the cosine similarity between *v_n_*(*X*) and *v_n_*_−1_(*X*), and vector magnitude was calculated using the Euclidean norm as a function of sequence length (***Supplementary Figs. S4***, ***S5*** and ***S7***).

For GFP-scaffold extensions, the same displacement analysis was applied to each scaffold precursor and its corresponding N-terminal or C-terminal extension. The resulting vectors were analyzed separately from the minimal de novo peptide vectors to quantify scaffold-dependent changes in direction consistency and magnitude (***Supplementary Figs. S2***, ***S3*** and ***S7***).

Differential vectors were calculated in the original high-dimensional representation space. Low-dimensional coordinates were used to display sequence states and connect trajectory points for visualization.

### Quantitative Metrics for Structural Space Analysis

Let *x*_1_, …, *x_N_* denote the embedding vectors in a local sequence ensemble. **Spatial Dispersion (*D*)** was defined as the mean pairwise Euclidean distance:

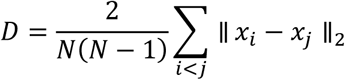

***D*** therefore quantifies the overall spread of an ensemble in representation space. For each point *x_i_*, we calculated the mean distance to its *k* nearest neighbours, where

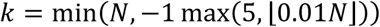

**Local Density (*ρ*)** was defined as

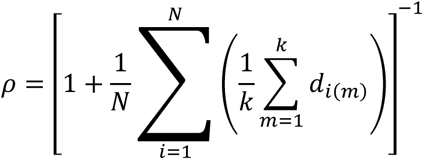

where *d_i_*_(_*_m_*_)_ denotes the Euclidean distance from *x_i_* to its *m* th nearest neighbour. This bounded definition gives 0 < *ρ* ≤ 1, with larger values indicating greater local compactness.

**Differential Embedding (*δ*)** was defined as

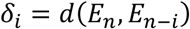

with the Manhattan distance used for the canonical sequential analysis. Euclidean distance was used to quantify complementary displacement magnitude, and cosine distance was defined as

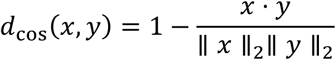

Accordingly, ***D*** describes ensemble spread, ***ρ*** describes local compactness and ***δ*** describes sequential displacement. These three metrics provide operational coordinates for identifying candidate structural anchors, perturbation-sensitive regions and abrupt state changes.

Because mean pooling can produce identical vectors for distinct single-site mutants, exact duplicate rows were quantified for each model and reported with the GFP mutation analyses. The effect of duplicate vectors on *ρ* must be evaluated using a prespecified sensitivity analysis and reported together with the primary density profile (***Supplementary Fig. S13***).

For sequence-elongation analysis, *δ* quantified displacement between successive sequence states, with the Manhattan norm used as the canonical measure. Cosine distance quantified directional reorientation. For local mutation or design ensembles, *D* and *ρ* quantified ensemble spread and compactness. For phase-transition analysis, Gaussian Wasserstein-2 distance quantified shifts in ensemble distribution by incorporating changes in both mean and covariance.

### Distributional Transition Analysis

A phase-transition-like representation-space event was defined as a distributional shift between consecutive sequence states. For this analysis, each model’s raw mean-pooled embedding matrix was independently mapped to a model-specific 32-dimensional random Gaussian projection space. Specifically, for model *m*, an embedding matrix 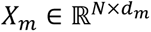 was multiplied by a projection matrix 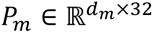, whose entries were sampled from 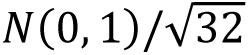 using random seed 42. The original embedding dimensions were *d_m_* = 1024 for ProstT5 and ProtT5, *d_m_* = 2560 for ESMC-6B and *d_m_* = 512 for METL-G. The projection was performed without centering or scaling, and each model therefore retained its own 32-dimensional projected representation space.

For each pair of adjacent sequence states, a Gaussian distribution was fitted to the corresponding projected ensemble. The squared Gaussian Wasserstein-2 distance was calculated as

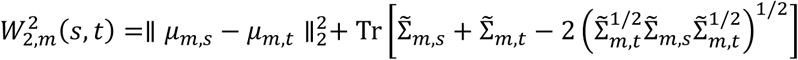

where *μ_m_*_,*s*_ and *μ_m_*_,*t*_ are the state-specific means and Σ̃*_m_*_,*s*_ and Σ̃*_m_*_,*t*_ are the regularized covariance matrices. To stabilize covariance estimation, Σ̃*_m_*_,*s*_ = Σ*_m_*_,*s*_ + *λ_m_I*_32_, with *λ_m_* = 10^−6^ for ProstT5 and *λ_m_* = 10^−8^ for ProtT5, ESMC-6B and METL-G. The implementation returned 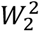 without the final square-root operation; accordingly, all raw values and standardized scores reported in this analysis refer to the squared Gaussian Wasserstein-2 distance.

Within each model, 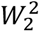 values were standardized into z-scores and accompanied by bootstrap 95% confidence intervals. ProstT5 confidence intervals were estimated using 200 bootstrap replicates, whereas ProtT5, ESMC-6B and METL-G used 500 replicates. CUSUM change-point detection was applied as a complementary analysis. Manhattan, Euclidean and cosine measures were used to characterize single-step displacement and directional reorientation, thereby providing complementary geometric descriptions of distributional state transitions.

### Protein Structure Prediction

For structural validation and high-resolution visualization, three-dimensional protein structures were predicted using locally installed implementations of *AlphaFold3*^14^ and *Boltz-2*^16^ with default parameters (Random seed with cycle = 10). To ensure topological consistency in *de novo* designed proteins, structural predictions were cross-validated using both *Boltz-2* and *AlphaFold3*. Model quality was assessed using the predicted Local Distance Difference Test (pLDDT) and predicted TM-score (pTM), where available. pLDDT values reported on a 0–1 scale were obtained by normalization of the original model scores.

Representative fixed-backbone ProteinMPNN designs were structurally evaluated using AlphaFold3 and cross-checked using Boltz-2. As a foldability-negative control, we generated 100 random sequences of length 239 and predicted their structures using AlphaFold3. These sequences had a mean normalized pLDDT of 0.413 ± 0.009, with no sequence exceeding 0.7 (***Supplementary Fig. S14***).

For the tTA-to-rtTA conformational comparison, we sampled BioEmu molecular conformational ensembles containing 500 conformations for each endpoint state, WT1 and ON1. These ensembles were analysed alongside the sequence-derived protein language model ensembles as complementary representations of state organization (***Supplementary Fig. S17***).

### Stepwise Genesis Simulation

To reconstruct protein genesis trajectories in silico, we performed systematic stepwise sequence elongation. For GFP and the TRIM-family proteins, we generated successive N-terminal prefixes of increasing length, beginning with a single residue and ending with the full-length protein. Each intermediate sequence was embedded using the protein language model and analysed as a discrete sequence state along a continuous representation-space trajectory. Critical features identified by *D*, *ρ* and *δ* were mapped onto AlphaFold3-predicted structures to examine their correspondence with predicted secondary-structure elements, domain boundaries and functional regions.

At the module scale, the 239-residue GFP was decomposed into 11 structurally coherent modules according to the AlphaFold3-predicted topology. Genesis was reconstructed using four assembly strategies: N-to-C, C-to-N, midpoint-outward and midpoint-asymmetric. For each accumulated module state, we calculated cosine-based step displacement, fragment Spatial Dispersion *D*, defined as the mean pairwise *L*_2_ distance among accumulated fragment representations, and fragment Local Density *ρ*.

The cosine-based step displacement was used to detect directional reorientation between consecutive module states, *D* was used to quantify the spread of the accumulated fragment ensemble and *ρ* was used to quantify its local compactness. This process-matched metric design enabled separation of module-level direction changes, ensemble expansion and local convergence.

### Mutational Landscape Analysis

To probe the microscopic organization of the protein representation space, we performed comprehensive single-site saturation mutagenesis across the full-length GFP sequence. We generated all 19 non-native amino acid substitutions at each of the 239 residue positions, yielding 4,541 single-site variants. For each position, the local mutation ensemble comprised the wild-type residue and its 19 non-native substitutions. These sequences were embedded and analysed using the mutation-indexed forms of Spatial Dispersion *D_i_*, Local Density *ρ_i_* and Differential Embedding *δ_i_* (***Supplementary Fig. S11***).

To assess cross-model consistency, mutation-associated profiles were generated using ProstT5, ProtT5, ESMC-6B, ESM2-650M and METL-G. Position-wise Spearman correlations were calculated between model-specific *δ_L_*_1_ profiles. Residue conservation was evaluated using 13 GFP homologues retrieved from Swiss-Prot by BLASTP with *E* ≤ 0.0001. After sequence alignment, the fraction of homologues retaining the GFP residue at each aligned position was calculated and converted into a percentile relative to the position-wise conservation background.

For the tTA-to-rtTA functional switch, we reconstructed the stepwise trajectory using six single-residue substitutions: S12G, E19G, A56P, F86Y, D148E and H179R. For each of the seven sequence states, we generated a single-mutation ensemble containing 3,915 sequences, yielding 27,405 sequences in total. These ensembles were embedded using ProstT5, ProtT5, ESMC-6B and METL-G and analysed using pointwise displacement metrics and Gaussian Wasserstein-2 distance.

The phase-transition analysis used squared Gaussian Wasserstein-2 distance because the transition was defined at the level of ensemble distribution. Manhattan displacement, Euclidean displacement and cosine distance were used as complementary measures of single-step movement and directional change. Within each model, *W*_2_ scores were standardized into z-scores, and bootstrap confidence intervals and CUSUM change-point analysis were applied as described above. The corresponding BioEmu conformational ensembles were analysed separately using the same endpoint and intermediate definitions.

### *De Novo* Design with ProteinMPNN

Fixed-backbone sequence design was performed using ProteinMPNN with GFP as the structural scaffold. We generated 2,000 sequences at each of four sampling temperatures, yielding 8,000 fixed-backbone designed variants in total. The exact temperature values and ProteinMPNN sampling parameters should be reported in the final Methods and Supplementary Methods. All 8,000 sequences were embedded and analyzed in the ProtGenesis representation space. For each temperature condition, the sequence with the highest prespecified ProteinMPNN recovery score was selected for structural validation using AlphaFold3. Representative designs were cross-checked using Boltz-2 where indicated.

The resulting design ensembles were compared with the human-proteome background in representation space. Sampling temperature *T* was treated as a controllable parameter for exploring the fixed-backbone design space. Spatial Dispersion *D*, Local Density *ρ* and Differential Embedding *δ* were calculated for the design ensembles, and PCA, t-SNE and UMAP were used for visualization. All quantitative metric values were calculated in the original high-dimensional representation space, whereas low-dimensional projections were used for display and trajectory illustration.

## Acknowledgments

This work was funded by the NUDT Research Program, the National Natural Science Foundation of China project (No. 32401056), the Hunan Province Science and Technology Innovation Program (2024RC3144, 2025JJ20028)

## Data and Code availability statement

The data and scripts used are saved in GitHub https://github.com/Travis13197/ProtGenesis.

## Ethics statement

Not applicable

## Supplementary Figures

**Fig. S1.**
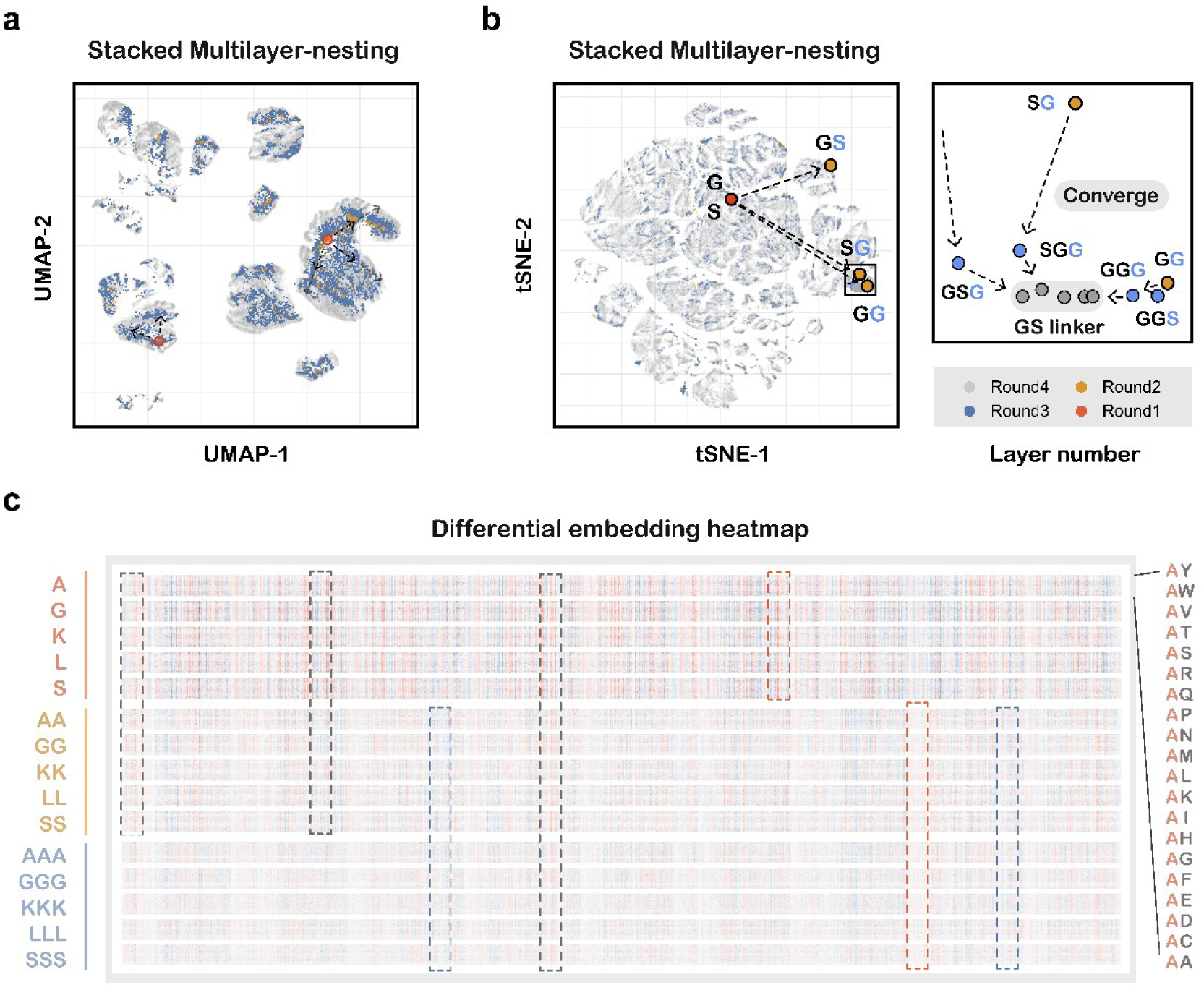
Hierarchical organization of the complete short-peptide representation space. **a,** Two-dimensional UMAP visualization of ProstT5 embeddings for all peptides of lengths 1–4, showing organization across sequence-length layers. **b,** t-SNE visualization of sequential peptide-generation rounds. **c,** Heatmap of 1,024-dimensional differential embeddings across hierarchical layers. These projections are used for visualization; quantitative distances were calculated in the original embedding space.

**Fig. S2.**
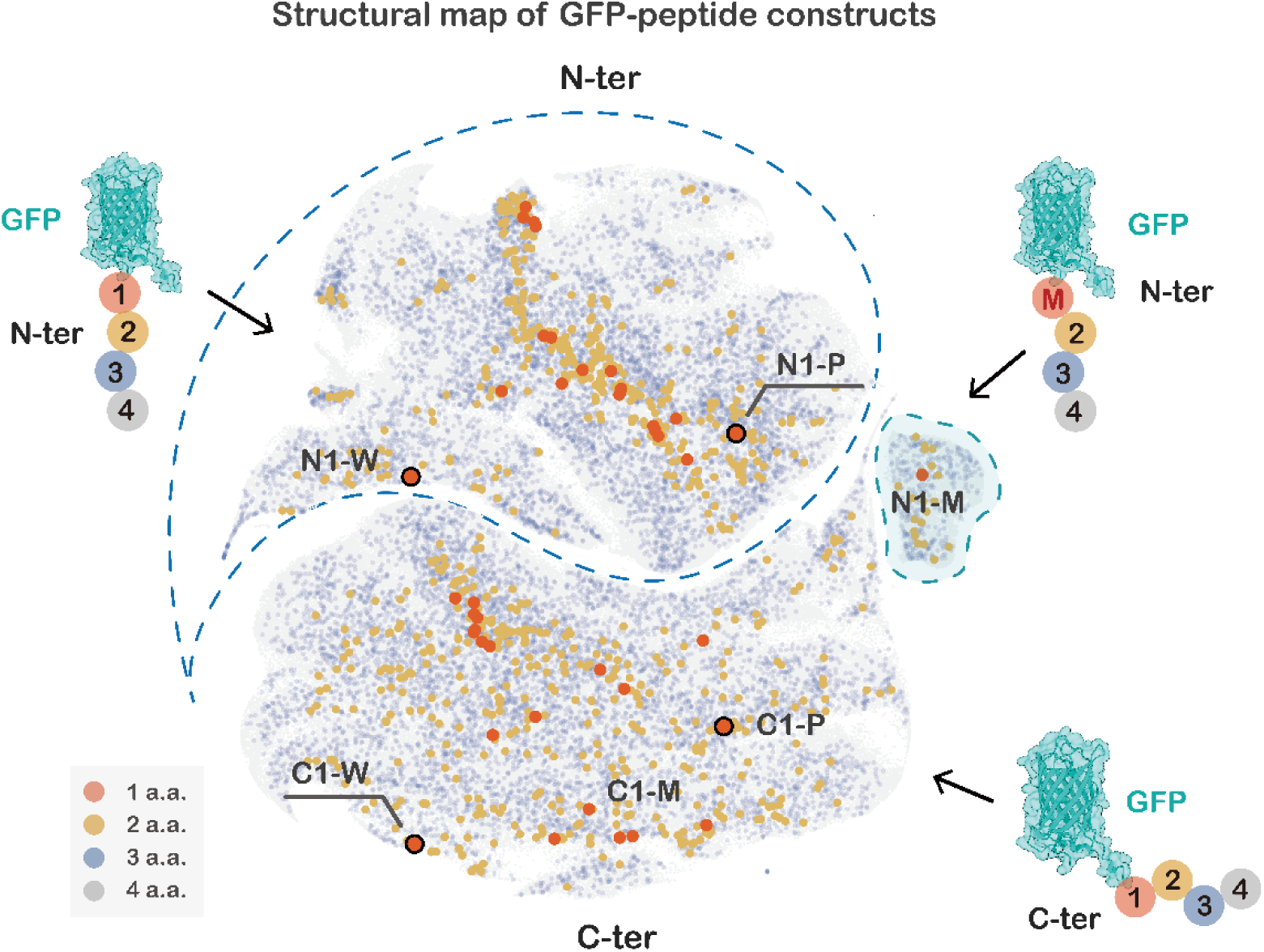
Terminal-context partitioning in GFP-scaffold extensions. Aggregate representation-space map of 336,840 GFP constructs containing all possible N-terminal or C-terminal extensions of one to four residues. The separation between N-terminal and C-terminal extension sets indicates scaffold- and terminal-context-dependent organization in representation space.

**Fig. S3.**
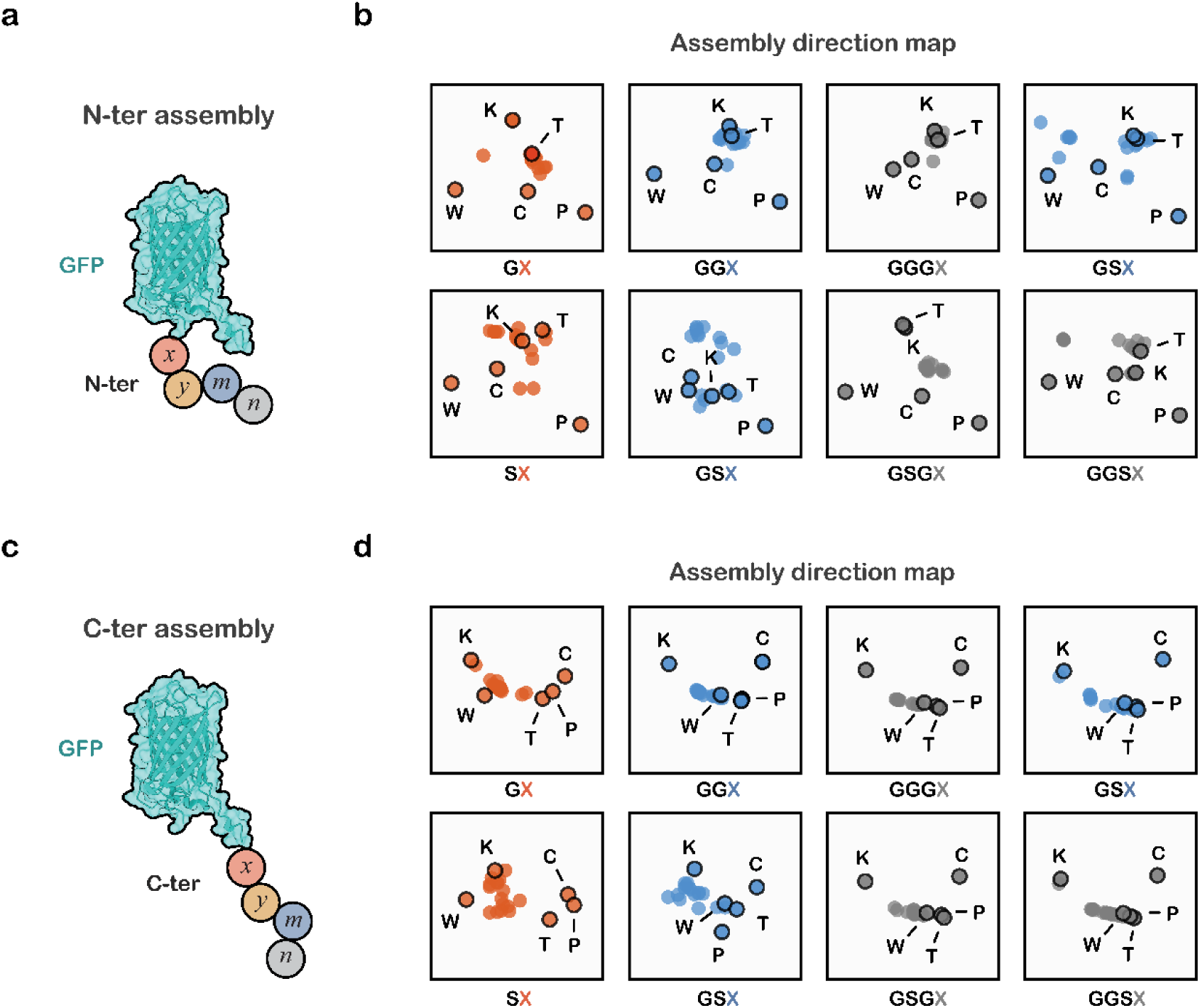
Differential embedding analysis of GFP-scaffold extensions. **a,c,** Construction of the N-terminal and C-terminal extension libraries. **b,d,** Assembly Direction Maps based on the displacement between each extended sequence and its corresponding GFP precursor. The maps illustrate context-dependent residue-associated displacement patterns in the scaffold regime.

**Fig. S4.**
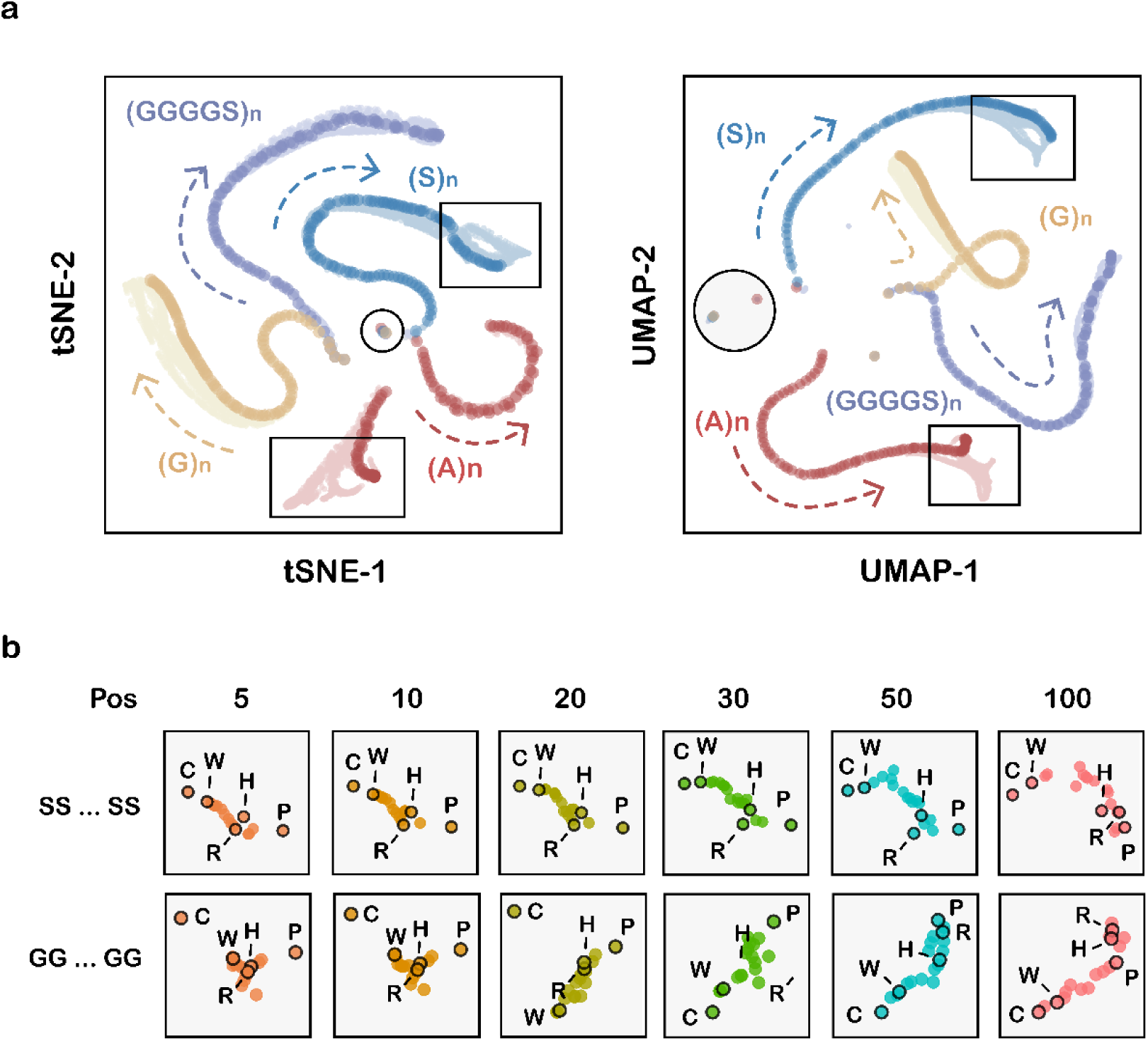
Representation-space trajectories in long low-complexity sequences. **a,** Construction of homopolymers and flexible linkers, including (*G*)*_n_*, (*S*)*_n_* and (*GGGGS*)*_n_*, with lengths of up to 120 residues. **b,** Assembly Direction Maps for sequential extension of representative low-complexity sequences. Residue-associated directional signatures remain structured but are progressively modulated by sequence length and context.

**Fig. S5.**
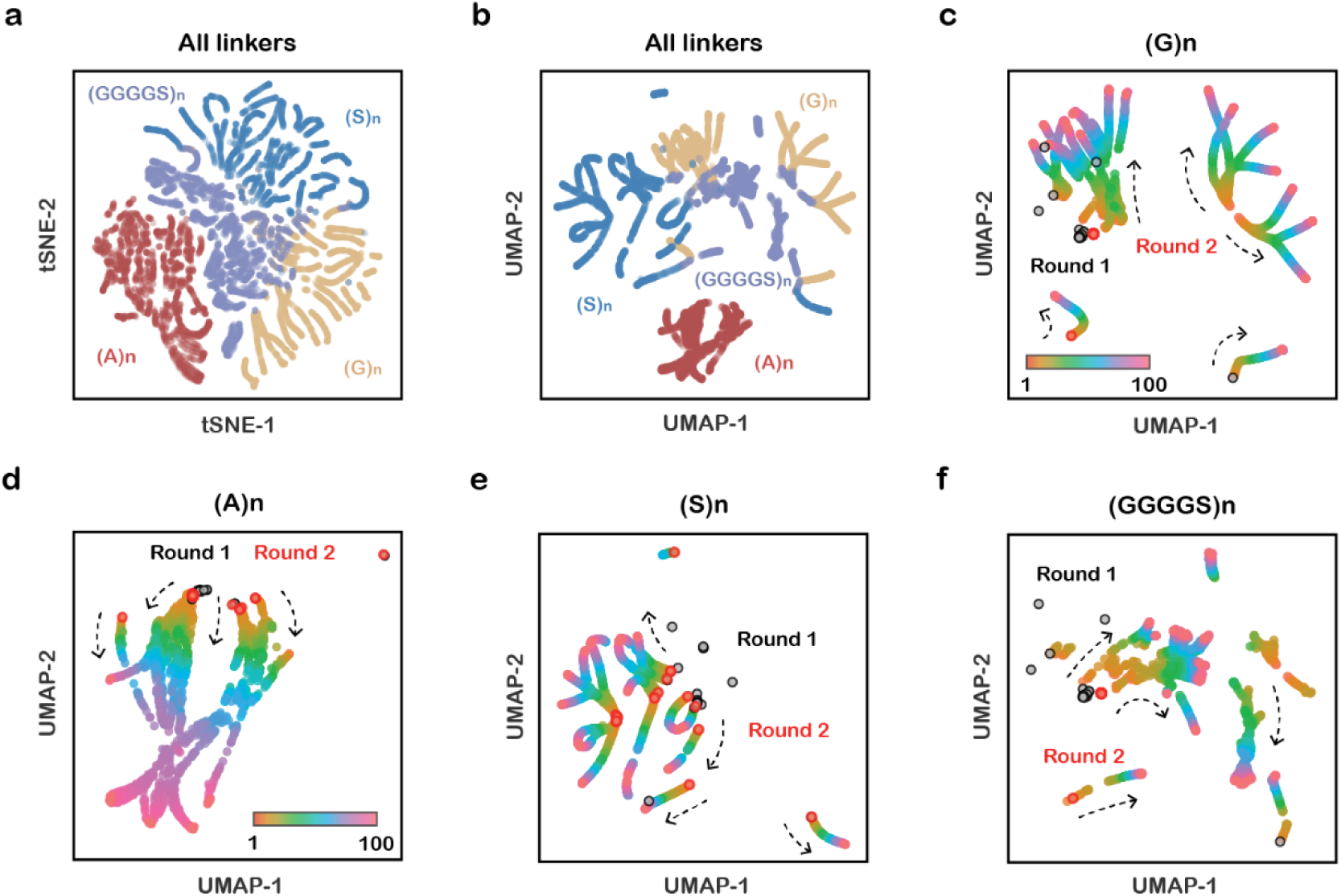
Position-dependent assembly trajectories in long homopolymers. **a,b,** Differential embedding analysis across successive positions in four homopolymeric sequences. **c–f,** Position-dependent Assembly Direction Maps. The results show reproducible but evolving representation-space trajectories as sequence context accumulates.

**Fig. S6.**
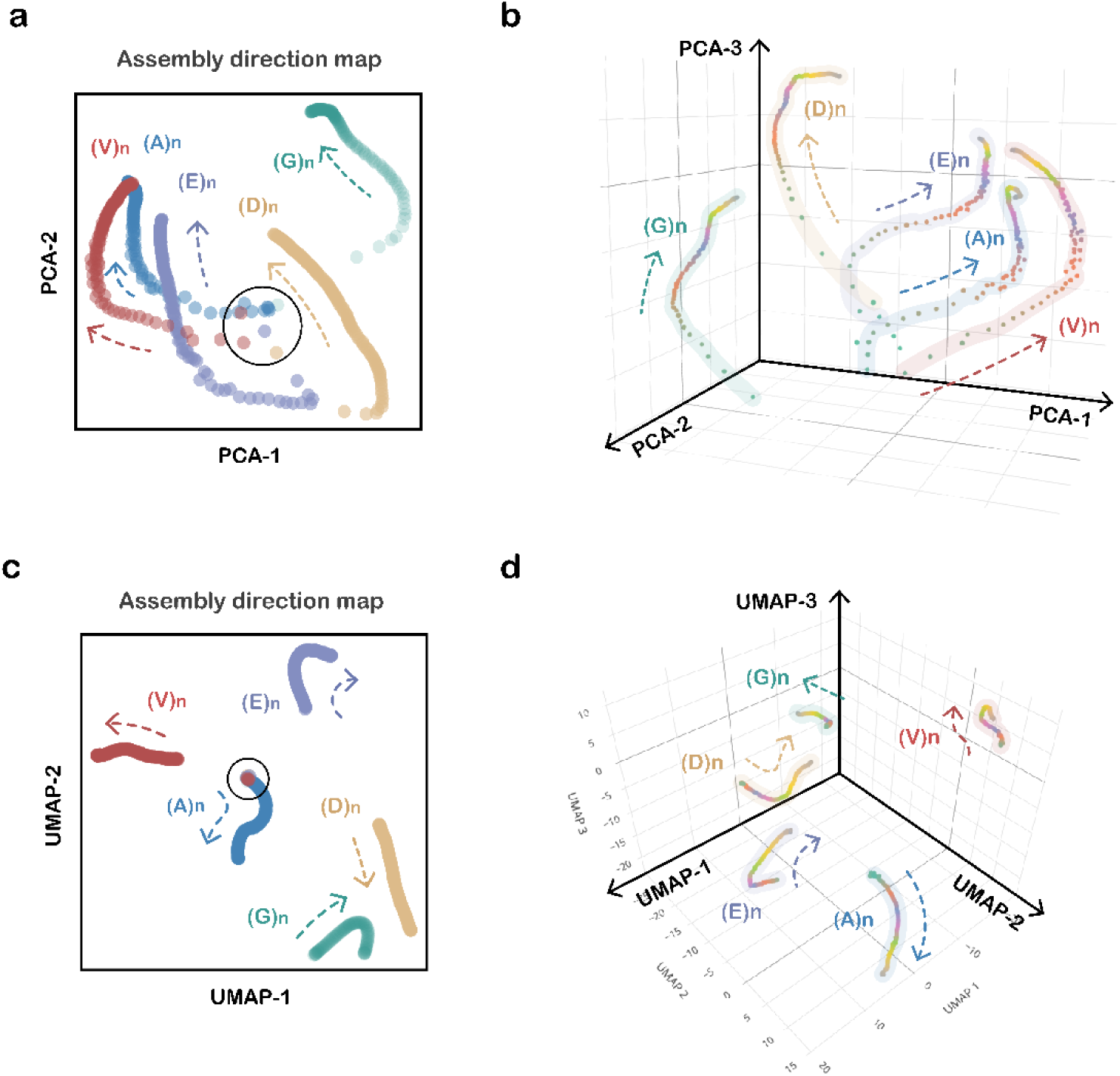
Amino-acid-associated trajectories in early-Earth-motivated homopolymers. **a,b,** Assembly Direction Maps for homopolymers derived from glycine, alanine, aspartate, valine and glutamate. **c,d,** Three-dimensional visualizations of the corresponding trajectories. These simplified sequences exhibit amino-acid-associated organization in representation space but do not directly reconstruct prebiotic polymerization or establish prebiotic physical laws.

**Fig. S7.**
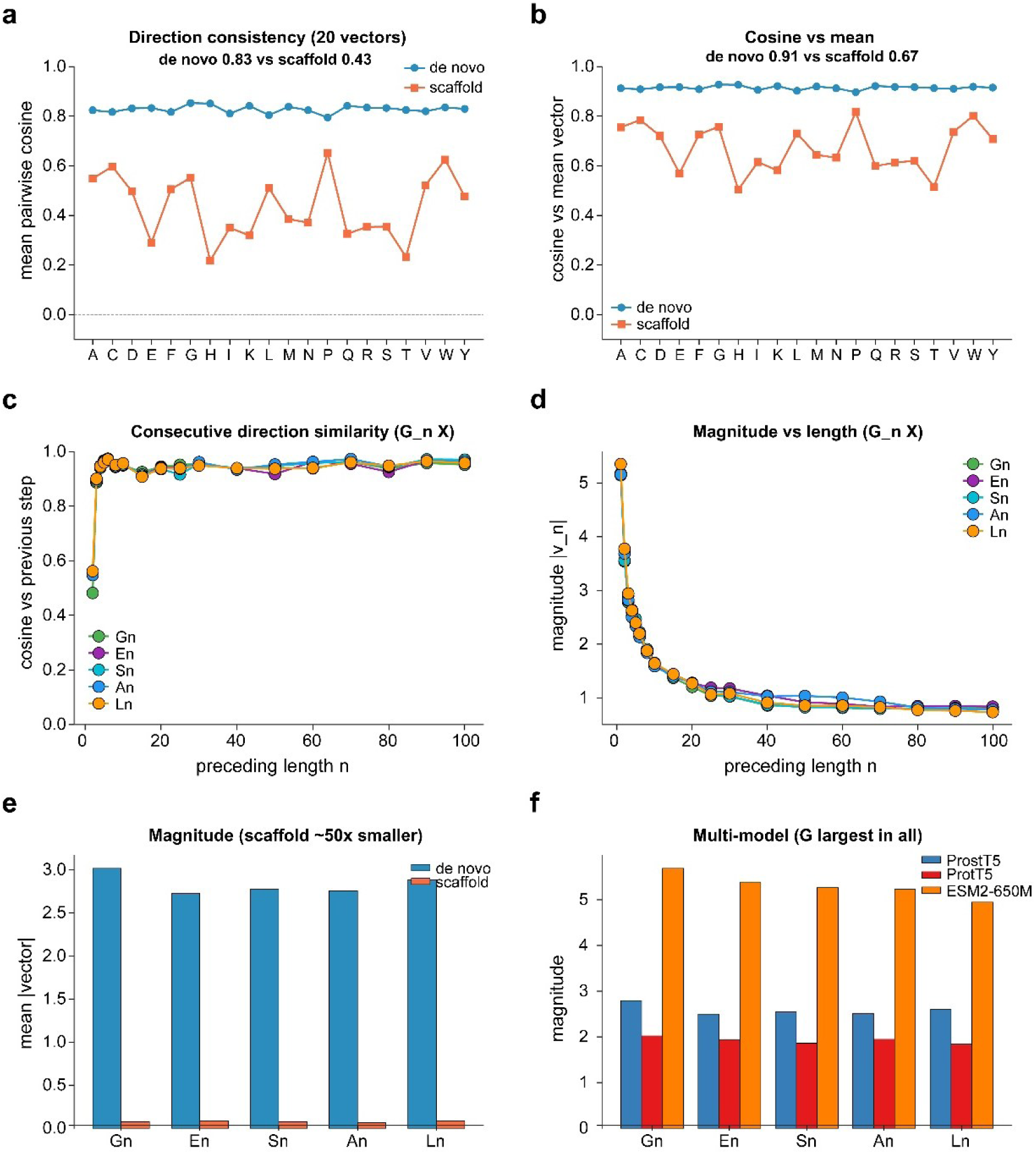
Assembly-vector context sensitivity. For each appended residue *X*, the differential vectors *v_a_*(*X*) = emb(*aX*) − emb(*a*), where *a* denotes one of the 20 preceding-residue contexts, quantify the representation-space displacement associated with adding *X*. In the minimal *de novo* regime, the mean pairwise cosine similarity among the 20 context-conditioned vectors was 0.83, and the mean cosine similarity relative to their average vector was 0.91. The corresponding values for GFP-scaffold extensions were 0.43 and 0.67, respectively. Across the tested *de novo* models, glycine showed the largest mean vector magnitude. These results indicate a reproducible but context-sensitive amino-acid assembly signature rather than a context-independent universal operator.

**Fig. S8.**
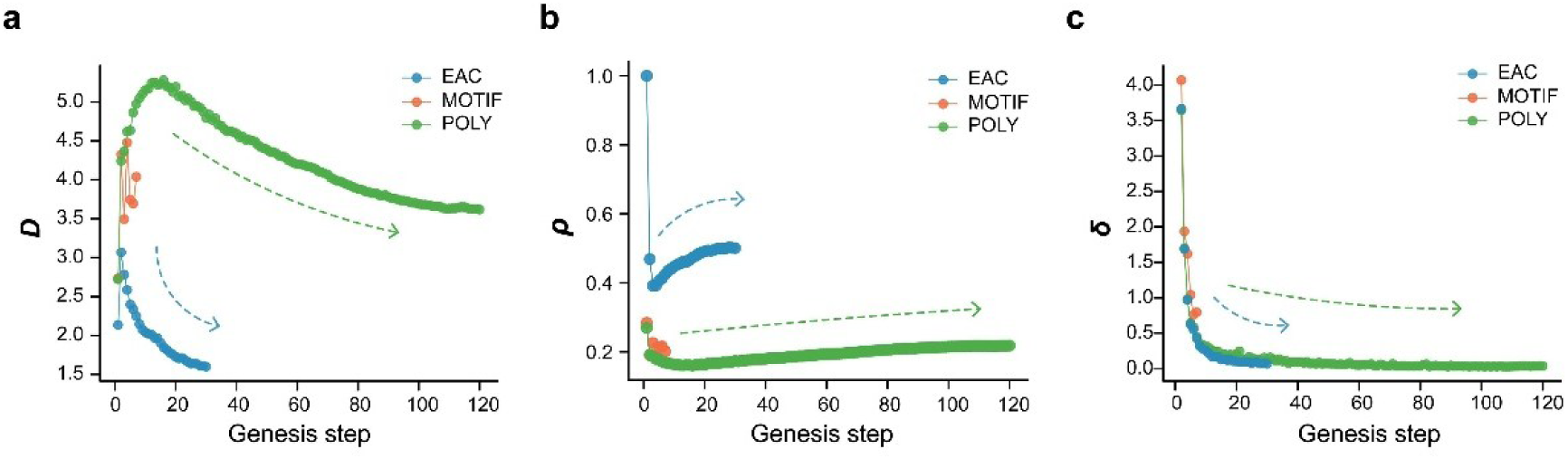
Genesis metric trends in early-Earth-motivated sequence cohorts. EAC denotes 100 combinatorial sequences generated from an early-amino-acid set and tracked for 30 steps; MOTIF denotes three short motifs tracked for seven steps; POLY denotes five homopolymers tracked for 120 steps. **a,** Dispersion *D* decreased with genesis step in the EAC cohort (*r_s_* = −0.986, *P* = 2 × 10^−2^^3^) and the POLY cohort (*r_s_* = −0.919, *P* = 2 × 10^−4^^9^). **b,** Local-density trends. **c,** Differential embedding distance across genesis steps. These results indicate progressive contraction of the representation-space ensemble and show that the strongest directional assembly signature is concentrated in the early short-range regime.

**Fig. S9.**
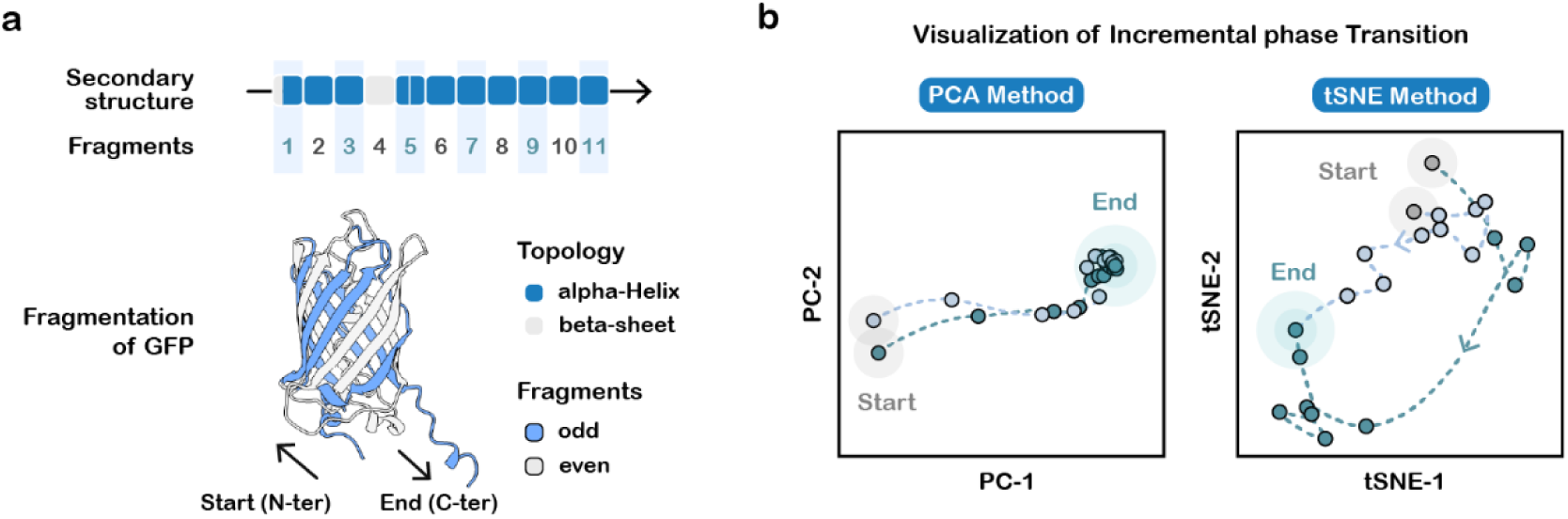
Incremental GFP module assembly generates discrete representation-space changes. **a,** Decomposition of GFP into 11 structurally coherent modules based on AlphaFold3-predicted topology. **b,** Representation-space trajectories generated by the N-to-C, C-to-N, midpoint-outward and midpoint-asymmetric assembly strategies. The trajectories contain gradual changes interspersed with larger stepwise displacements.

**Fig. S10.**
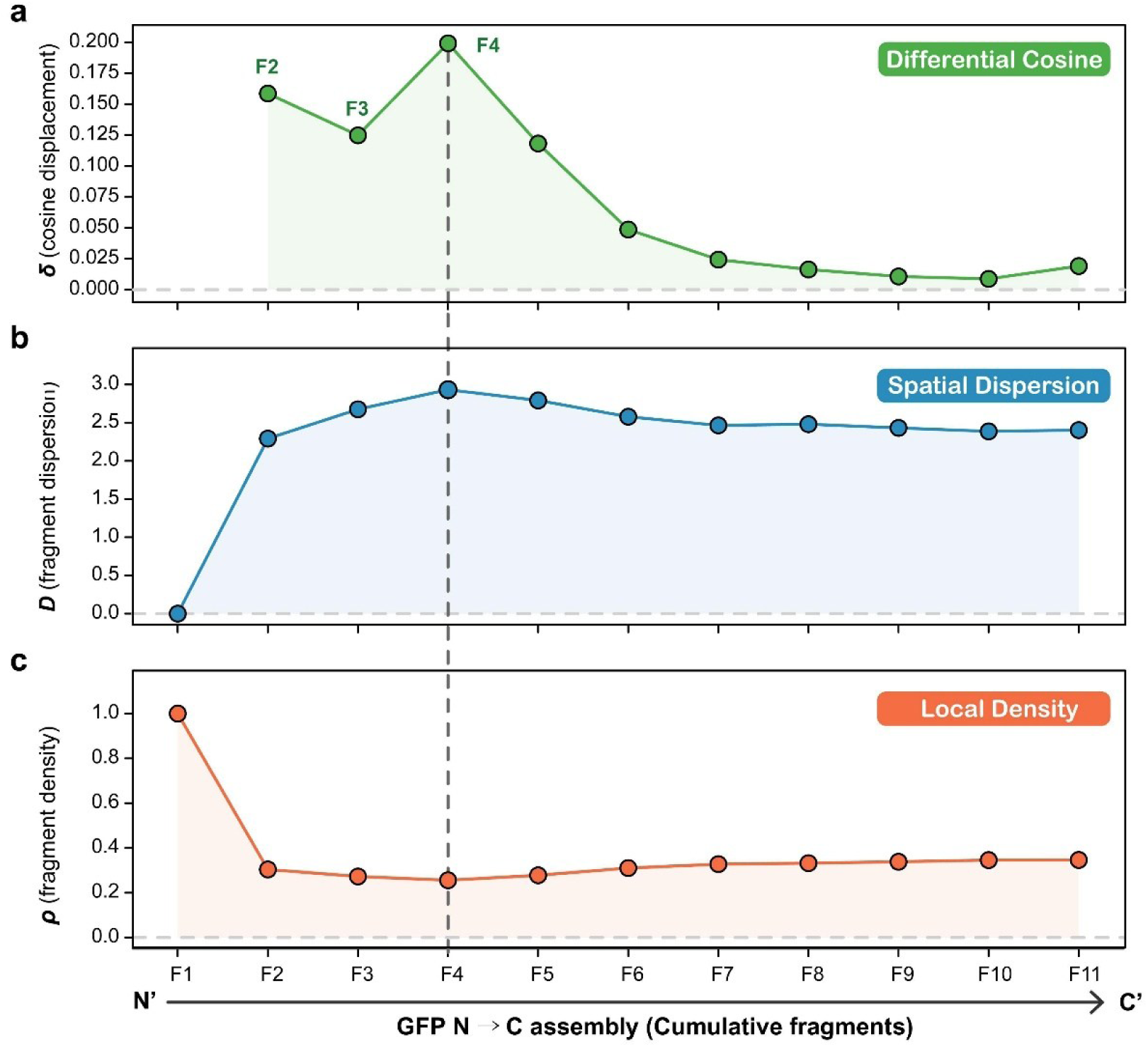
Multi-metric analysis of the GFP N-to-C module-assembly trajectory. **a,** Cosine-based step displacement, with prominent peaks at F2 and F4. **b,** Fragment Spatial Dispersion *D*, reaching a maximum at F4 (*D* = 2.93), the chromophore-containing fragment. **c,** Fragment Local Density *ρ*, which decreases monotonically across the assembly sequence.

**Fig. S11.**
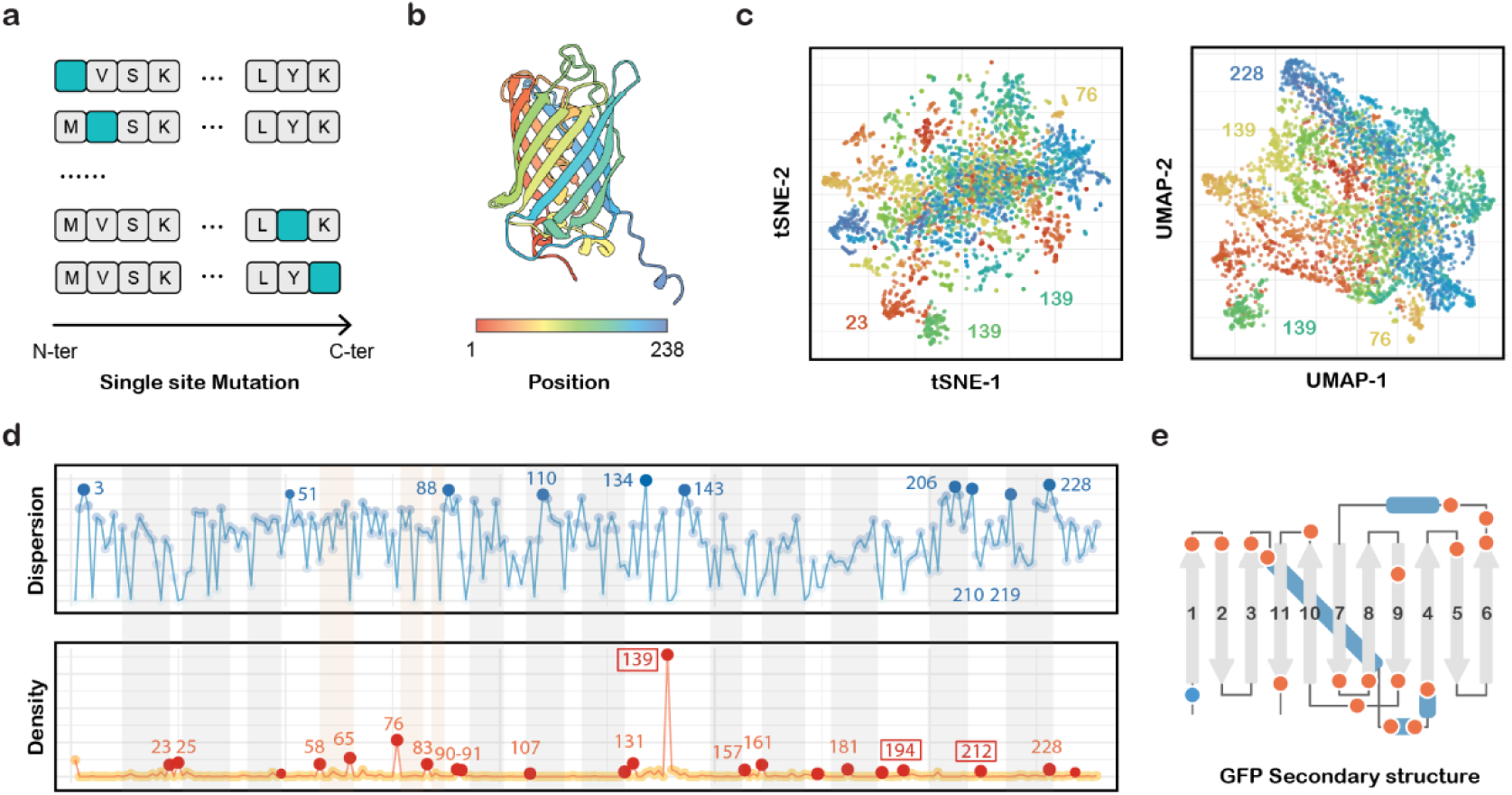
Full-length single-site mutation landscape of GFP. **a,b,** Representation-space analysis of the complete single-site substitution ensemble. **c,** t-SNE and UMAP visualizations. **d,e,** Mutation-indexed dispersion, local-density and differential-distance profiles mapped onto GFP secondary-structure elements. The metrics describe representation-space perturbation geometry rather than direct fitness or thermodynamic stability.

**Fig. S12.**
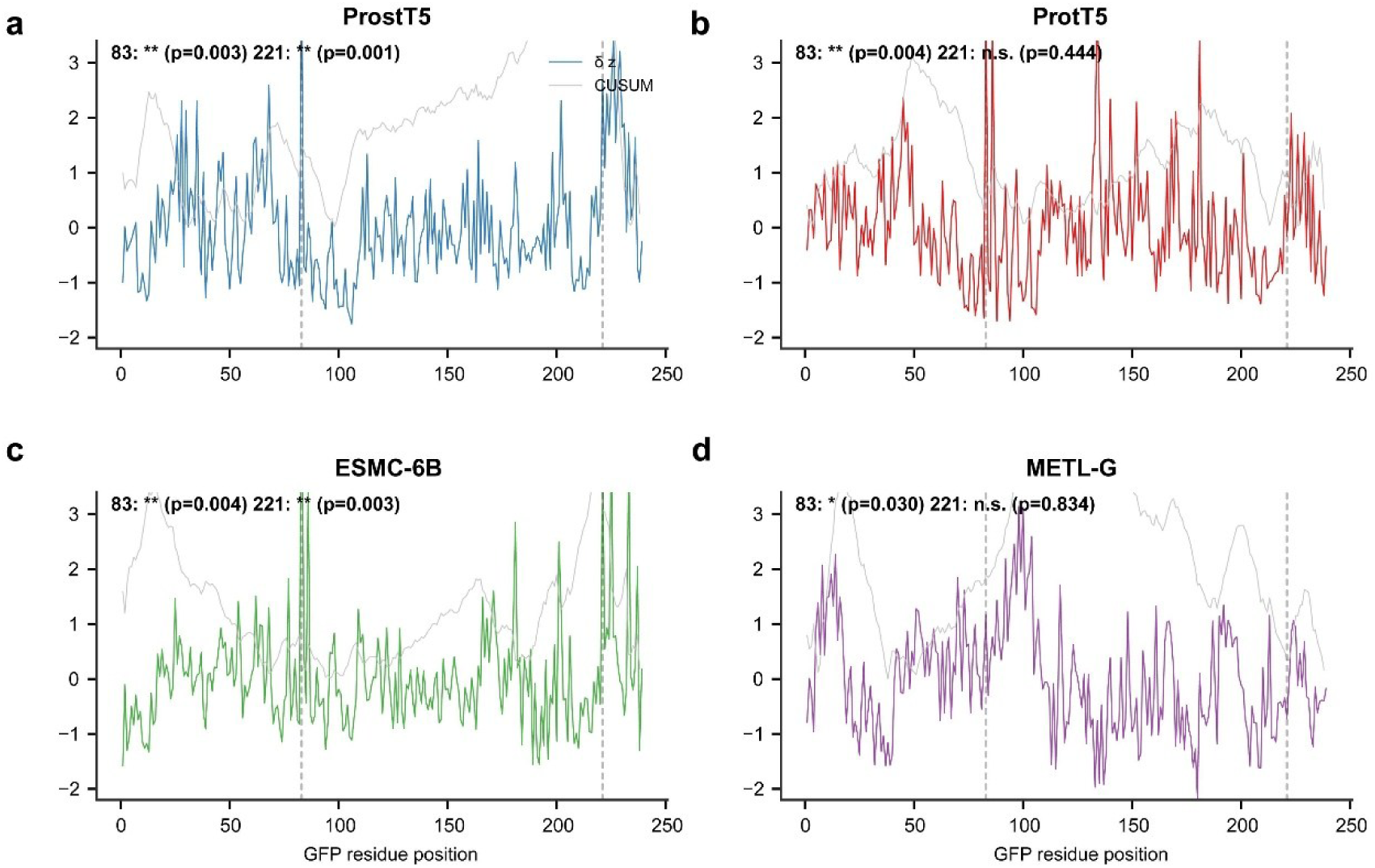
Four-model GFP single-mutation analysis. Mutation-associated representation-space profiles generated using ProstT5, ProtT5, ESMC-6B and METL-G. Residue 83 was identified in four models and residue 221 in three models when the complete five-model analysis was considered.

**Fig. S13.**
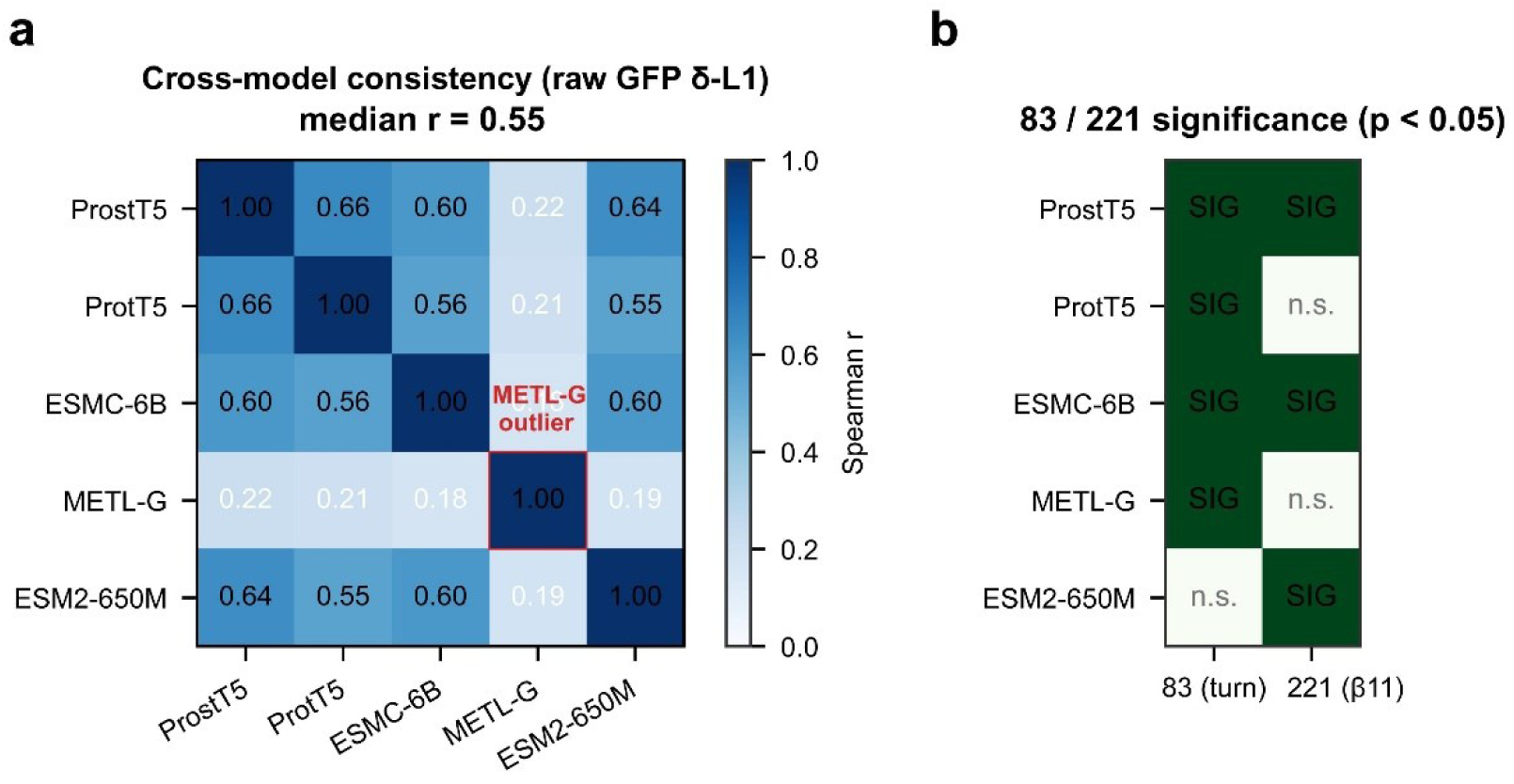
Five-model consistency of GFP mutation-associated representation features. ProstT5, ProtT5, ESMC-6B and ESM2-650M showed moderate agreement in raw *δ* profiles (*r_s_* = 0.55– 0.66), whereas METL-G was less concordant with the other models (*r_s_* = 0.18-0.22). Residue 83 was significant in four of five models, and residue 221 was significant in three of five models. Exact duplicate rows generated by mean pooling are reported and considered when interpreting density-based metrics.

**Fig. S14.**
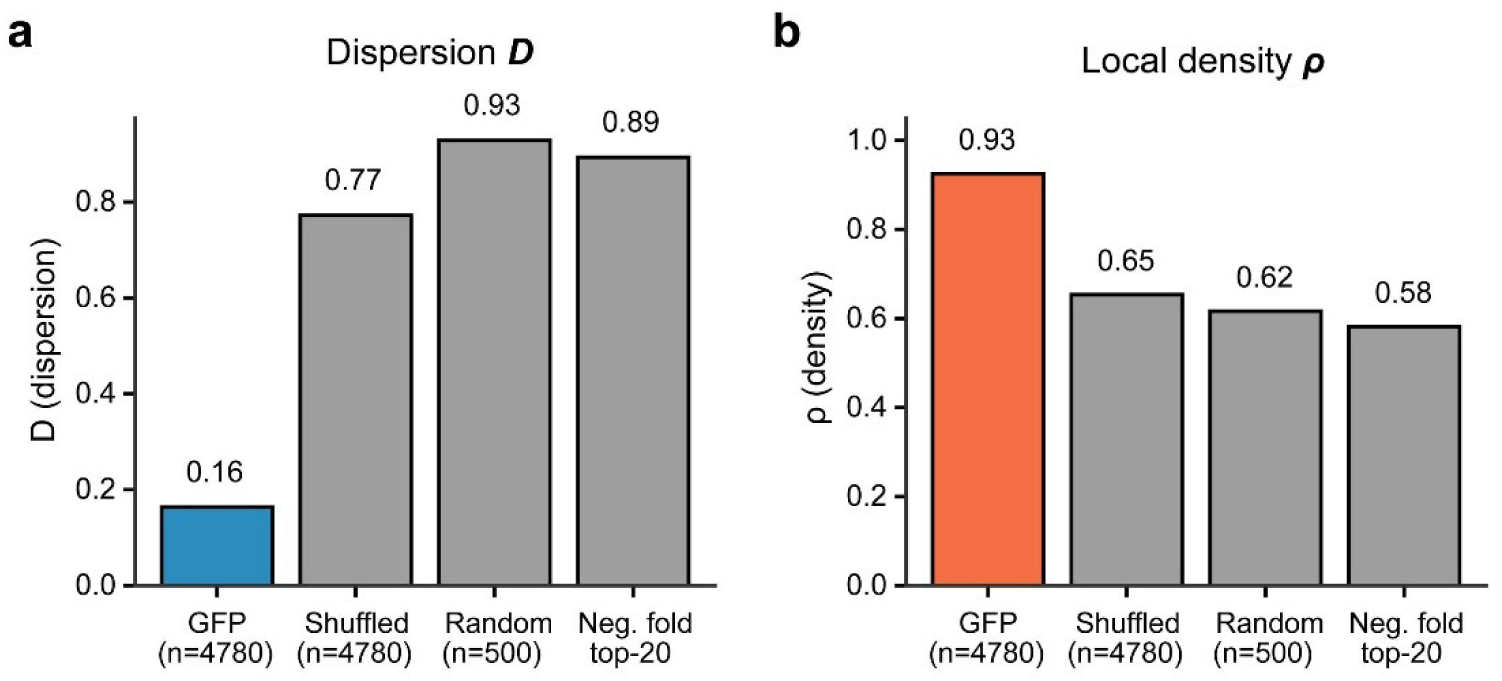
Foldability-negative control. One hundred random 239-residue sequences had a mean normalized pLDDT of 0.413 ± 0.009, and none exceeded 0.7. Their representation-space metrics differed from those of the GFP ensemble, with *D* = 0.89 and *ρ* = 0.58, compared with *D* = 0.16 and *ρ* = 0.93 for GFP.

**Fig. S15.**
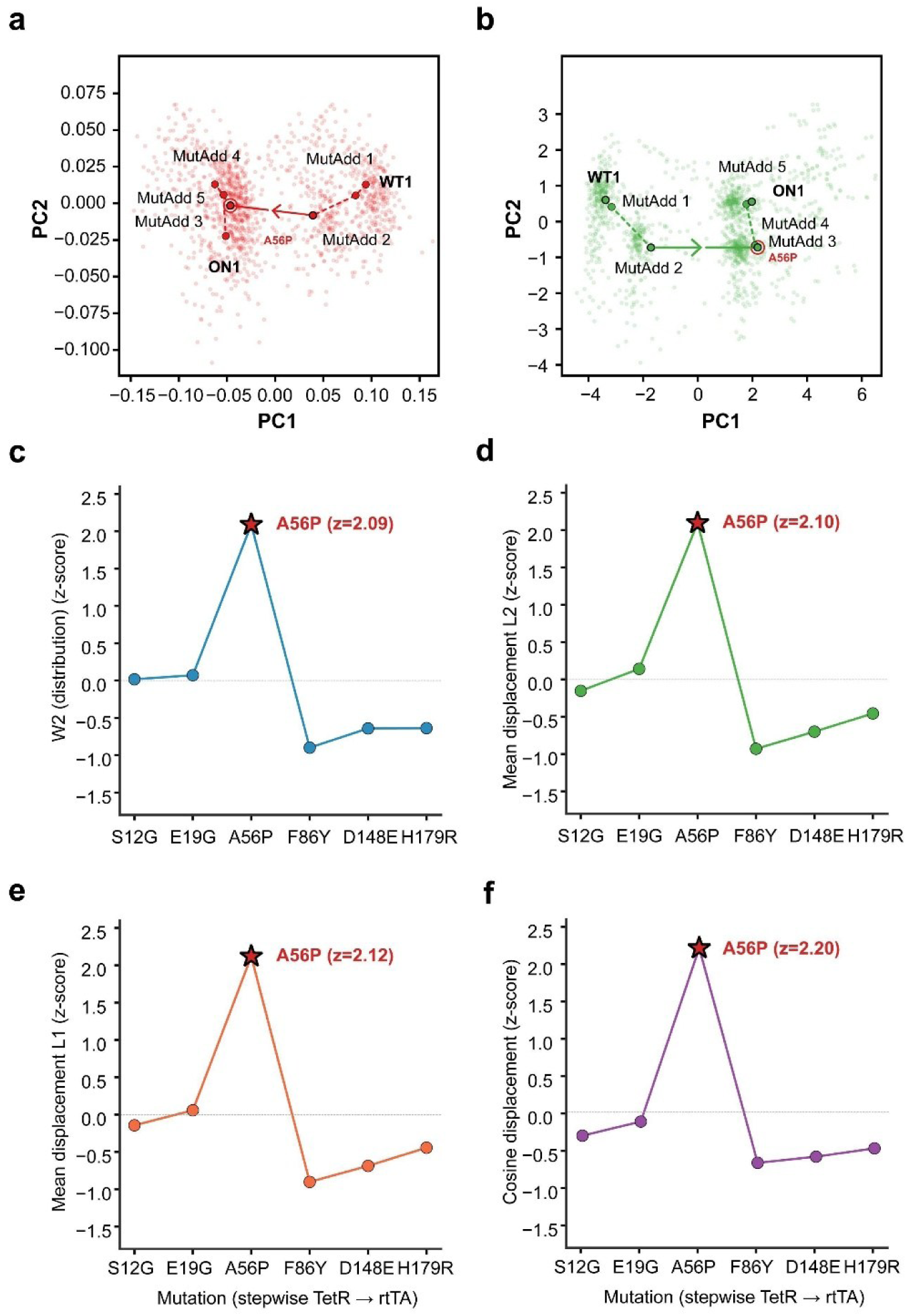
Four-model analysis of the tTA-to-rtTA sequence trajectory. **a,b,** PCA visualization of state centroids, identifying A56P as the largest stepwise displacement. **c–f,** ProstT5-based multi-metric analysis using squared Gaussian Wasserstein-2 distance (*W*^2^), mean-displacement *L*_2_, mean-displacement *L*_1_and cosine displacement. A56P, corresponding to the MutAdd_2_→MutAdd_3_ step, was the maximal transition under all four measures in ProstT5. The corresponding within-model Wasserstein-2 z-scores were 2.09, 2.17, 2.21 and 2.09 across ProstT5, ProtT5, ESMC-6B and METL-G.

**Fig. S16.**
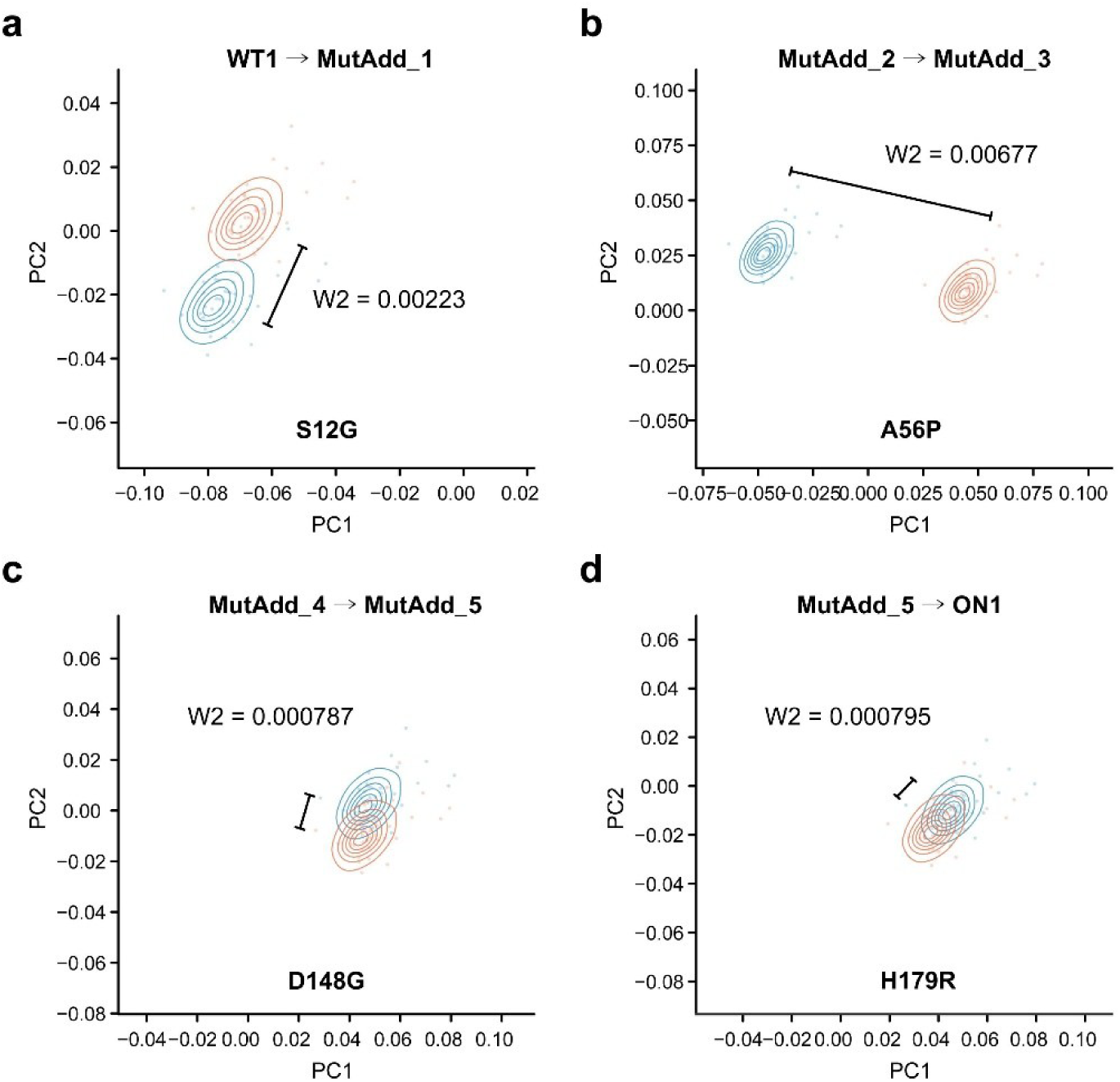
Single-mutation ensemble distributions along the tTA-to-rtTA trajectory. Two-dimensional density contours for selected consecutive state transitions, each generated from 3,915 single-mutation sequences per state and annotated with the corresponding squared Gaussian Wasserstein-2 value (*W*^2^). The largest distributional change occurs at the A56P-associated transition.

**Fig. S17.**
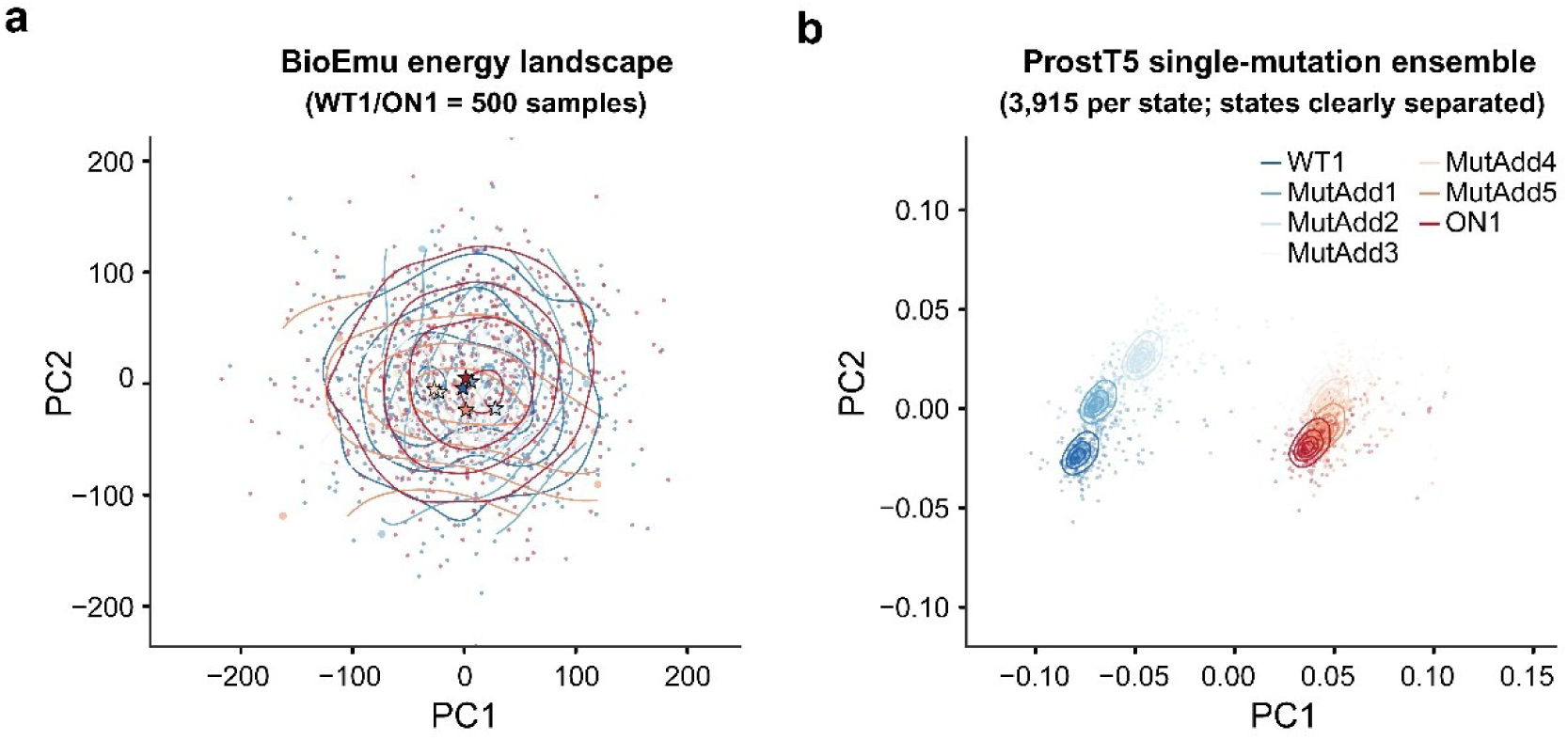
Comparison of sequence-derived representation ensembles with BioEmu conformational ensembles. **a,** BioEmu conformational ensembles containing 500 sampled conformations for the WT1 and ON1 endpoint states. **b,** Protein language model single-mutation ensembles containing 3,915 sequences per state. The two analyses sample different objects - molecular conformations and sequence-derived representation states - and are therefore shown as complementary rather than as direct measurements of a common energy landscape.

**Fig. S18.**
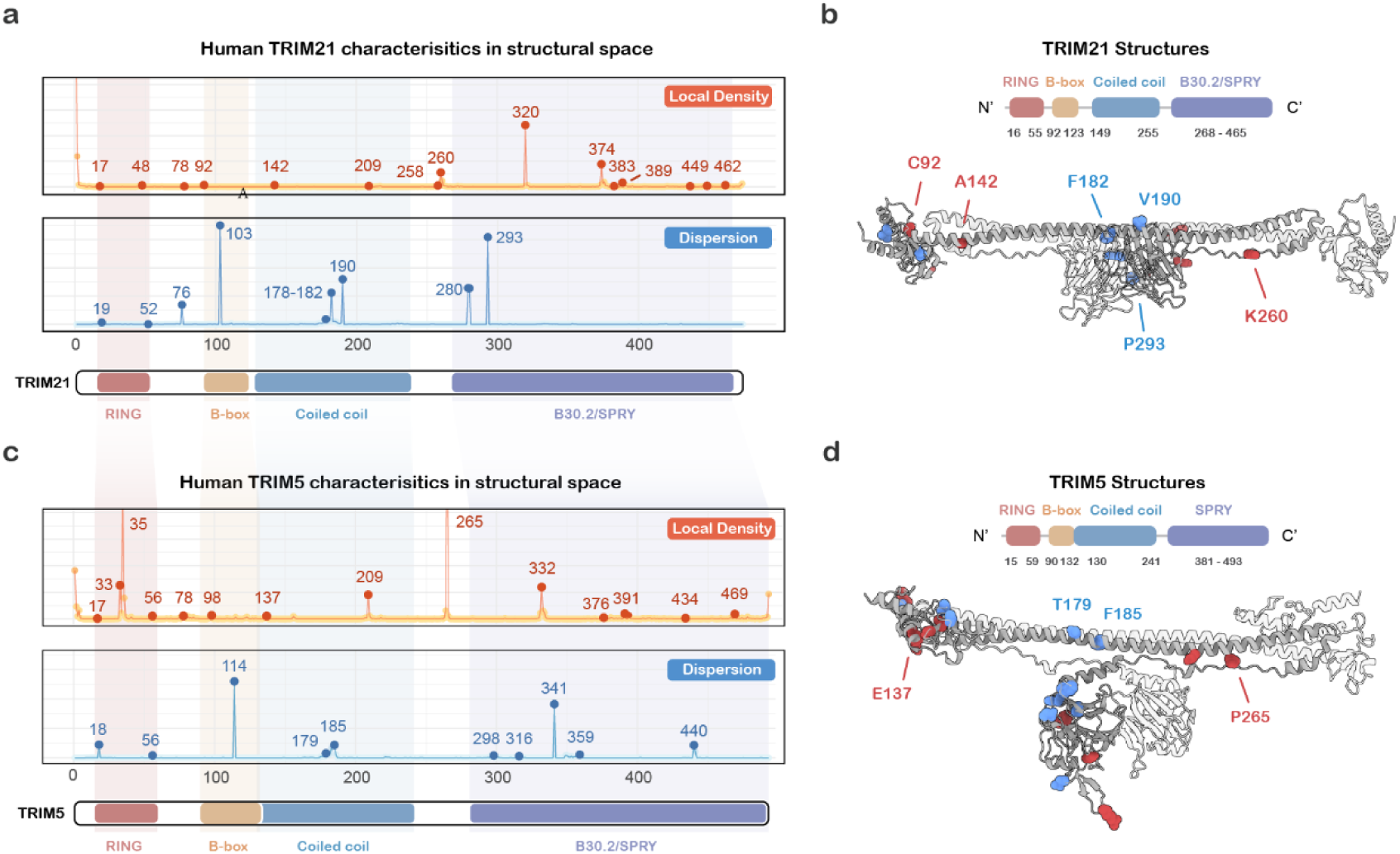
Stepwise representation-space trajectories of TRIM-family E3 ligases. **a,c,** Sequential N-terminal-prefix trajectories of human TRIM21 and TRIM5. **b,d,** Domain architectures and metric-associated features mapped onto AlphaFold3-predicted structures. Recurring density- and dispersion-associated features coincide with selected modular or functional landmarks in both proteins.

**Fig. S19.**
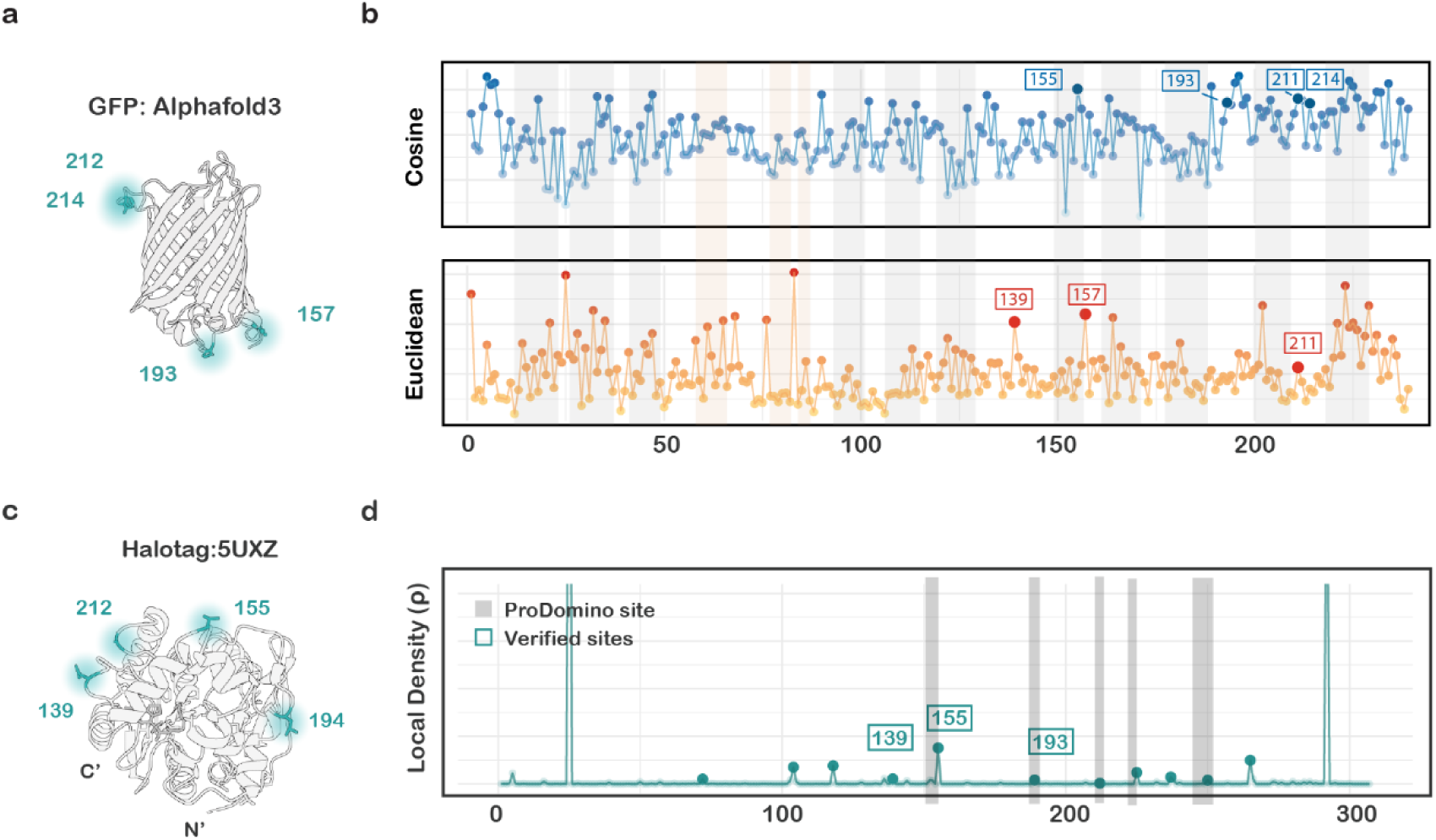
Retrospective comparison of representation-space features with validated split-protein sites. **a,** Experimentally reported GFP split sites mapped onto the predicted GFP structure. **b,** Spatial-distance and local-density analysis of GFP single-site variants. **c,** Structural mapping of a reported HaloTag split site. **d,** Local-density profile of HaloTag. The correspondence between density-associated features and reported sites supports candidate-site prioritization but does not establish prospective predictive performance.

**Fig. S20.**
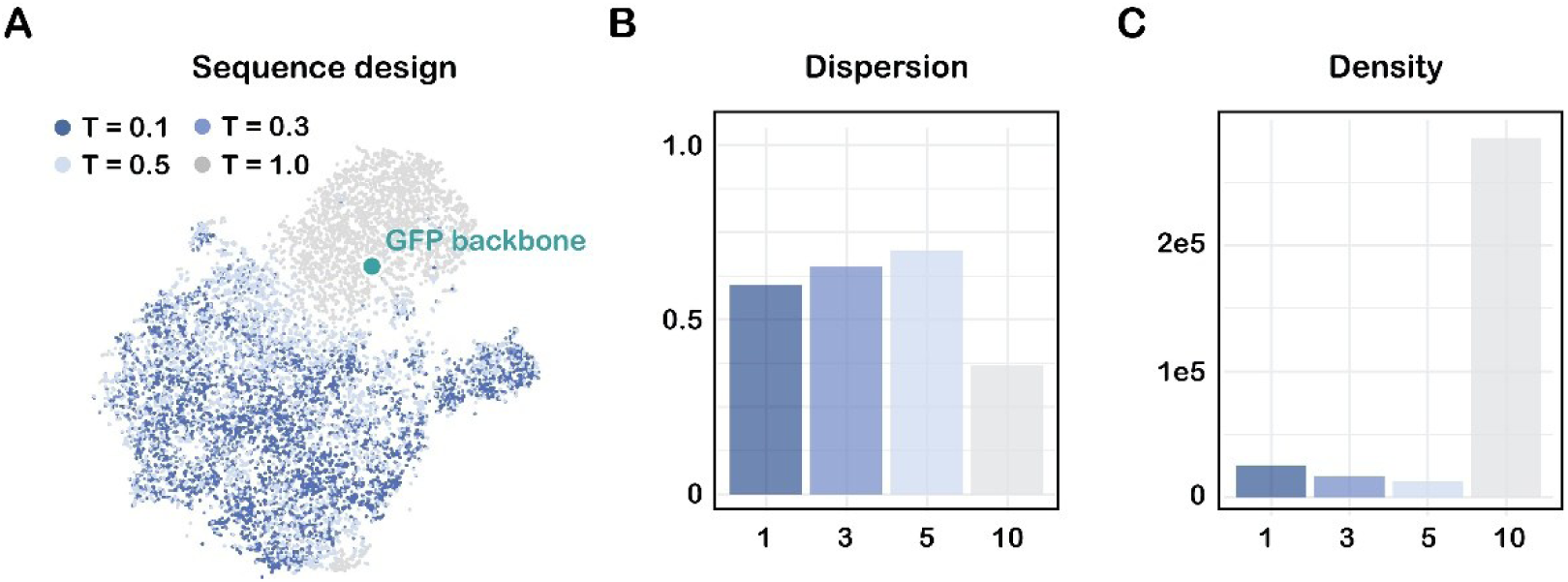
ProteinMPNN fixed-backbone design ensembles across sampling temperatures. **a,** t-SNE visualization of 8,000 GFP fixed-backbone designed variants generated at four sampling temperatures. **b,c,** Spatial Dispersion *D* and Local Density *ρ* across design conditions. The metrics quantify temperature-associated redistribution and expansion of the design ensemble in representation space. The analyses describe controllable exploration of a fixed-backbone design space and should not be interpreted as direct evidence of fully de novo structural generation.

## Reference

1 Kortemme, T. De novo protein design-From new structures to programmable functions. Cell 187, 526–544 (2024). 10.1016/j.cell.2023.12.028

2 Notin, P., Rollins, N., Gal, Y., Sander, C. & Marks, D. Machine learning for functional protein design. Nat Biotechnol 42, 216–228 (2024). 10.1038/s41587-024-02127-0

3 Huang, P. S., Boyken, S. E. & Baker, D. The coming of age of de novo protein design. Nature 537, 320–327 (2016). 10.1038/nature19946

4 Bunne, C. et al. How to build the virtual cell with artificial intelligence: Priorities and opportunities. Cell 187, 7045–7063 (2024). 10.1016/j.cell.2024.11.015

5 Qian, L., Dong, Z. & Guo, T. Grow AI virtual cells: three data pillars and closed-loop learning. Cell Res 35, 319–321 (2025). 10.1038/s41422-025-01101-y

6 Higgs, P. G. & Lehman, N. The RNA World: molecular cooperation at the origins of life. Nat Rev Genet 16, 7–17 (2015). 10.1038/nrg3841

7 Zhang, B. & Cech, T. R. Peptide bond formation by in vitro selected ribozymes. Nature 390, 96–100 (1997). 10.1038/36375

8 Miller, S. L. A production of amino acids under possible primitive earth conditions. Science 117, 528–529 (1953). 10.1126/science.117.3046.528

9 Longo, L. M. et al. Primordial emergence of a nucleic acid-binding protein via phase separation and statistical ornithine-to-arginine conversion. Proc Natl Acad Sci U S A 117, 15731–15739 (2020). 10.1073/pnas.2001989117

10 Anfinsen, C. B. Principles that Govern the Folding of Protein Chains. Science 181, 223–230 (1973). 10.1126/science.181.4096.223

11 Sahakyan, H., Babajanyan, S. G., Wolf, Y. I. & Koonin, E. V. In silico evolution of globular protein folds from random sequences. Proc Natl Acad Sci U S A 122, e2509015122 (2025). 10.1073/pnas.2509015122

12 Draizen, E. J., Veretnik, S., Mura, C. & Bourne, P. E. Deep generative models of protein structure uncover distant relationships across a continuous fold space. Nat Commun 15, 8094 (2024). 10.1038/s41467-024-52020-2

13 Rufo, C. M. et al. Short peptides self-assemble to produce catalytic amyloids. Nat Chem 6, 303–309 (2014). 10.1038/nchem.1894

14 Abramson, J. et al. Accurate structure prediction of biomolecular interactions with AlphaFold 3. Nature 630, 493–500 (2024). 10.1038/s41586-024-07487-w

15 Wohlwend, J. et al. Boltz-1 Democratizing Biomolecular Interaction Modeling. bioRxiv (2025). 10.1101/2024.11.19.624167

16 Passaro, S. et al. Boltz-2: Towards Accurate and Efficient Binding Affinity Prediction. bioRxiv (2025). 10.1101/2025.06.14.659707

17 Varadi, M. et al. AlphaFold Protein Structure Database in 2024: providing structure coverage for over 214 million protein sequences. Nucleic Acids Res 52, D368–d375 (2024). 10.1093/nar/gkad1011

18 Durairaj, J. et al. Uncovering new families and folds in the natural protein universe. Nature 622, 646–653 (2023). 10.1038/s41586-023-06622-3

19 Holm, L. & Sander, C. Mapping the protein universe. Science 273, 595–603 (1996). 10.1126/science.273.5275.595

20 Koonin, E. V., Wolf, Y. I. & Karev, G. P. The structure of the protein universe and genome evolution. Nature 420, 218–223 (2002). 10.1038/nature01256

21 Bileschi, M. L. et al. Using deep learning to annotate the protein universe. Nat Biotechnol 40, 932–937 (2022). 10.1038/s41587-021-01179-w

22 van Kempen, M. et al. Fast and accurate protein structure search with Foldseek. Nat Biotechnol 42, 243–246 (2024). 10.1038/s41587-023-01773-0

23 Barrio-Hernandez, I. et al. Clustering predicted structures at the scale of the known protein universe. Nature 622, 637–645 (2023). 10.1038/s41586-023-06510-w

24 Lau, A. M. et al. Exploring structural diversity across the protein universe with The Encyclopedia of Domains. Science 386, eadq4946 (2024). 10.1126/science.adq4946

25 Zheng, W. et al. Deep-learning-based single-domain and multidomain protein structure prediction with D-I-TASSER. Nat Biotechnol (2025). 10.1038/s41587-025-02654-4

26 Vázquez Torres, S., et al. De novo designed proteins neutralize lethal snake venom toxins. Nature 639, 225–231 (2025). 10.1038/s41586-024-08393-x

27 Lauko, A. et al. Computational design of serine hydrolases. Science 388, eadu2454 (2025). 10.1126/science.adu2454

28 Zhu, J. et al. De novo design of transmembrane fluorescence-activating proteins. Nature 640, 249–257 (2025). 10.1038/s41586-025-08598-8

29 Yu, B. et al. De novo design of light-responsive protein-protein interactions enables reversible formation of protein assemblies. Nat Chem (2025). 10.1038/s41557-025-01929-2

30 Braberg, H., Echeverria, I., Kaake, R. M., Sali, A. & Krogan, N. J. From systems to structure — using genetic data to model protein structures. Nature Reviews Genetics 23, 342–354 (2022). 10.1038/s41576-021-00441-w

31 Heinzinger, M. et al. Bilingual language model for protein sequence and structure. NAR Genom Bioinform 6, lqae150 (2024). 10.1093/nargab/lqae150

32 Elnaggar, A. et al. ProtTrans: Toward Understanding the Language of Life Through Self-Supervised Learning. IEEE Trans Pattern Anal Mach Intell 44, 7112–7127 (2022). 10.1109/tpami.2021.3095381

33 Tan, Y., et al. VenusX: Unlocking Fine-Grained Functional Understanding of Proteins. arXiv *preprint arXiv:2505.11812* (2025).

34 Hayes, T. et al. Simulating 500 million years of evolution with a language model. Science 387, 850–858 (2025). 10.1126/science.ads0018

35 Heinzinger, M. et al. ProstT5: Bilingual Language Model for Protein Sequence and Structure. bioRxiv, 2023.2007.2023.550085 (2023). 10.1101/2023.07.23.550085

36 Sangster, A. G. et al. Zero-shot segmentation using embeddings from a protein language model identifies functional regions in the human proteome. bioRxiv, 2025.2003.2005.641584 (2025). 10.1101/2025.03.05.641584

37 Zhang, G., Liu, C., Lu, J., Zhang, S. & Zhu, L. The Role of AI-Driven De Novo Protein Design in the Exploration of the Protein Functional Universe. Biology 14, 1268 (2025).

38 Gelman, S. et al. Biophysics-based protein language models for protein engineering. Nature Methods 22, 1868–1879 (2025). 10.1038/s41592-025-02776-2

39 Lewis, S. et al. Scalable emulation of protein equilibrium ensembles with generative deep learning. Science 389, eadv9817 (2025). 10.1126/science.adv9817

40 Clift, D. et al. A Method for the Acute and Rapid Degradation of Endogenous Proteins. Cell 171, 1692–1706.e1618 (2017). 10.1016/j.cell.2017.10.033

41 Zeng, J. et al. Target-induced clustering activates Trim-Away of pathogens and proteins. Nat Struct Mol Biol 28, 278–289 (2021). 10.1038/s41594-021-00560-2

42 Liu, C. et al. Predator: A novel method for targeted protein degradation. bioRxiv, 2020.2007.2031.231787 (2020). 10.1101/2020.07.31.231787

43 Bae, J., Kim, J., Choi, J., Lee, H. & Koh, M. Split Proteins and Reassembly Modules for Biological Applications. ChemBioChem 25, e202400123 (2024). 10.1002/cbic.202400123

44 Wolf, B. et al. Rational engineering of allosteric protein switches by in silico prediction of domain insertion sites. Nat Methods 22, 1698–1706 (2025). 10.1038/s41592-025-02741-z

45 Dauparas, J. et al. Robust deep learning-based protein sequence design using ProteinMPNN. Science 378, 49–56 (2022). 10.1126/science.add2187

46 Butcher, J. et al. De novo Design of All-atom Biomolecular Interactions with RFdiffusion3. bioRxiv (2025). 10.1101/2025.09.18.676967

47 Hartl, B., Levin, M. & Pio-Lopez, L. Neural cellular automata: Applications to biology and beyond classical AI. Phys Life Rev 56, 94–108 (2025). 10.1016/j.plrev.2025.11.010

48 Qiu, X., Zhu, L., Wang, H. & Xie, M. Biocomputing at the crossroad between emulating artificial intelligence and cellular supremacy. Curr. Opin. Biotechnol. 92, 103264 (2025). 10.1016/j.copbio.2025.103264

49 Candido, S. et al. Language Modeling Materializes a World Model of Protein Biology. bioRxiv, 2026.2006.2003.729735 (2026). 10.64898/2026.06.03.729735

